# Mapping the molecular and structural specialization of the skin basement membrane for inter-tissue interactions

**DOI:** 10.1101/2020.04.27.061952

**Authors:** Ko Tsutsui, Hiroki Machida, Ritsuko Morita, Asako Nakagawa, Kiyotoshi Sekiguchi, Jeffrey H. Miner, Hironobu Fujiwara

## Abstract

Inter-tissue interaction is fundamental to multicellularity. Although the basement membrane (BM) is located at tissue interfaces, its mode of action in inter-tissue interactions remains poorly understood, mainly because the molecular and structural details of the BM at distinct inter-tissue interfaces remain unclear. By combining quantitative transcriptomics and immunohistochemistry, we systematically identify the cellular origin, molecular identity and tissue distribution of extracellular matrix molecules in mouse hair follicles, and reveal that BM composition and architecture are exquisitely specialized for distinct inter-tissue interactions, including epidermal–fibroblast, epidermal–muscle and epidermal–nerve interactions. The epidermal–fibroblast interface, namely, hair germ–dermal papilla interface, makes asymmetrically organized side-specific heterogeneity in BM, defined by the newly characterized interface, hook and mesh BMs. One component of these BMs, laminin α5, is required for the topological and functional integrity of hair germ–dermal papilla interactions. Our study highlights the significance of BM heterogeneity in distinct inter-tissue interactions.

## Introduction

The extracellular matrix (ECM) is a complex noncellular network of multicellular organisms that plays essential roles in animal development and homeostasis. The basement membrane (BM) is a thin sheet-like ECM that is located at the border of tissues, where it serves to compartmentalize and also tightly integrate tissues^1, 2^. The BM has several crucial roles: i) it provides structural support to cells that is essential for the development of organ structures; ii) it signals to cells through adhesion receptors such as integrins; iii) it controls the tissue distributions and activities of soluble growth factors; and iv) its mechanical characteristics influence cell behavior^3, 4, 5^. Thus, the composition and structure of the BM play critical roles in cell proliferation, differentiation, migration, survival, polarity and positioning, underpinning many fundamental biological phenomena, including the developmental patterning, inter-tissue interactions and stem cell niche formation.

The BM is composed of a large variety of molecules that exhibit spatiotemporal expression patterns during development and homeostasis, indicating that individual cell types are exposed to tailor-made BM niches^6, 7, 8^. In mammals, the entire set of ECM molecules, called the matrixome or matrisome, is encoded by approximately 300 ECM genes and there are also approximately 800 ECM-associated genes, such as those encoding ECM-modifying enzymes and growth factors^7, 9^ (http://matrisomeproject.mit.edu/). Although information about the unique distribution, biochemical activities and *in vivo* functions of individual BM molecules has been accumulated, the entire molecular landscape of the BM composition, including its cellular origins, tissue localizations and pattern-forming processes, in all organs remains largely unknown. One major reason for the difficulty in obtaining a comprehensive understanding of the ECM’s composition lies in the biochemical properties of ECM proteins, including their large size, insolubility and crosslinked nature. These properties have made it challenging to systematically characterize ECM specialization in its entirety at the cellular resolution in tissues. This lack of knowledge impedes our understanding of the extrinsic regulation of cellular behaviour and interactions.

Mouse hair follicle (HF) is an excellent model to investigate the formation and function of spatiotemporally specialized ECMs because this mini-organ is tiny, yet has clear epidermal and dermal compartments that are associated with specific tissue architecture and functions (Fig. 1a)^10, 11^. Different types of epidermal stem cells are compartmentalized along the longitudinal axis of the HF and attach to the BM. These different epidermal stem cells are associated with distinct dermal cells, such as the lanceolate mechanosensory complexes in the upper bulge for touch sensation and epidermal stem cell regulation^12, 13, 14^, arrector pili muscle in the mid-bulge for piloerection^15^, and the dermal papilla (DP) in the hair germ (HG) for HF development and regeneration^16^. These epidermal–dermal interactions take place via the BM.

**Fig. 1.**
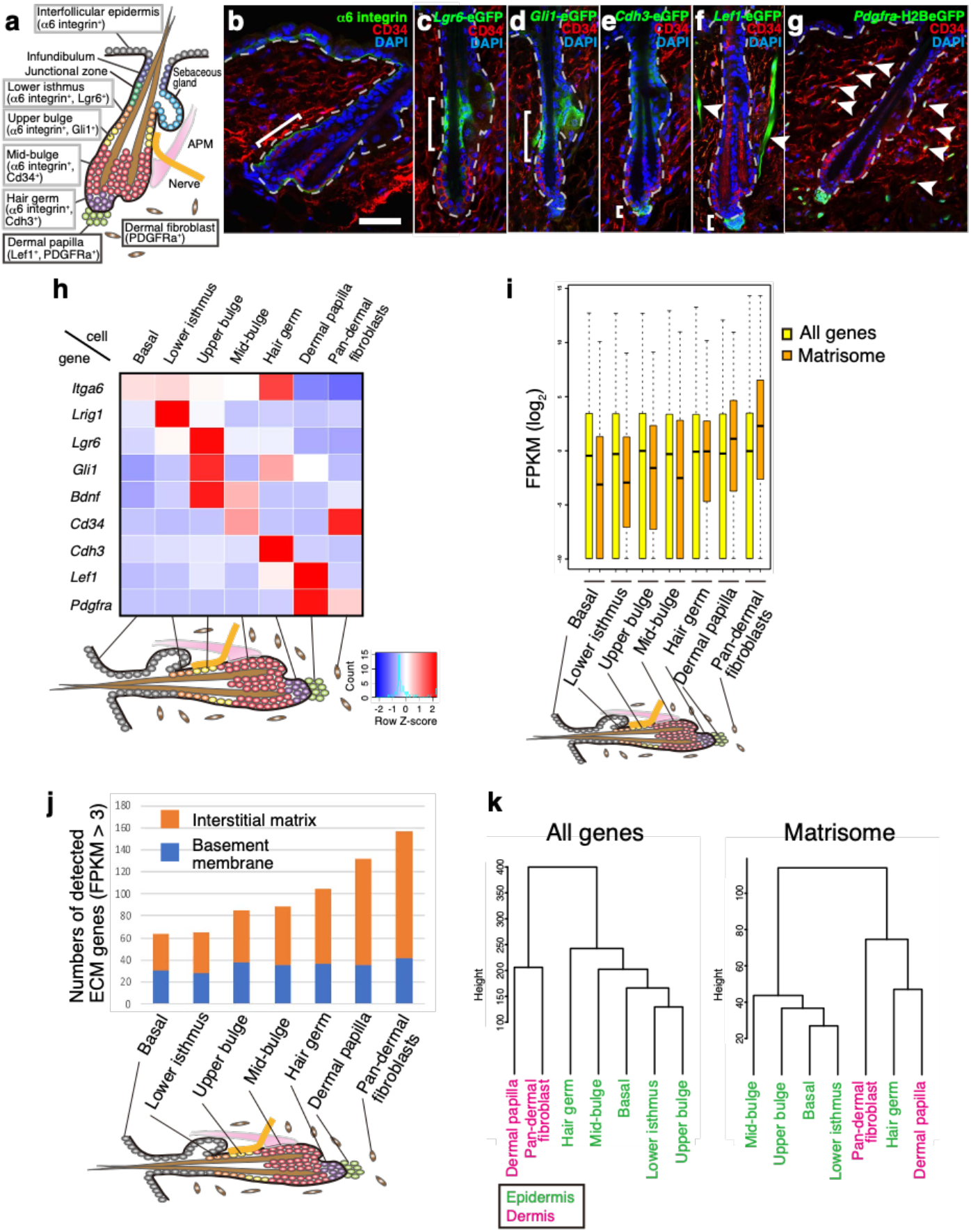
Targeted cell isolation and transcriptional profiling of the mouse hair follicle. **a** Graphical illustration of adult telogen HF compartments. Grey and black frames indicate epidermal and dermal compartments targeted in this study, respectively. APM, arrector pili muscle. **b-g** Tissue distribution of the markers for each HF compartment. Brackets indicate the target cell compartment in each panel. HF mid-bulge basal stem cells were labelled with α6 integrin (green) and CD34 (red) (b). Lower isthmus epidermal basal stem cells were visualized by *Lgr6*-eGFP (green) (c). Upper bulge epidermal basal stem cells were visualized by *Gli1*-eGFP (green) (d). HG cells were visualized by *Cdh3*-eGFP (green) (e). DP cells and arrector pili muscles (arrowheads) were visualized by *Lefl*-eGFP (green) (f). Dermal fibroblasts (arrowheads), including DP cells, were visualized by *Pdgfra*-H2BeGFP (green) (g). White dashed lines indicate epidermal-dermal borders. Scale bar, 50 μm. **h** Relative mRNA expression levels of HF region-specific genes in different sorted cell populations. mRNA levels are expressed relative to *Gapdh* and represented by Z-score values, **i** Expression levels of all genes and matrisome genes in different HF cell populations. Data are median FPKM (log_2_) with first to third quartile box and dashed line ranging from the minimum to the maximum. n=3 (HG and DP) or 4 (other populations). **j** Number of ECM genes detected in different HF cell populations. Genes with FPKM values greater than 3 were counted. Averaged FPKM values from three or four biological replicates were examined. **k** Global ECM gene expression correlation among HF cell populations examined by hierarchical clustering. Genes with FPKM values greater than 3 were used for the analysis among all genes (left panel) and matrisome genes (right panel). Averaged FPKM values from three or four biological replicates were examined. Green and magenta colours indicate epidermal and dermal compartments, respectively.

Previous studies suggested that the BM is an important niche component for both epidermal stem cells and dermal cells. Loss of contact with the ECM or reduced integrin expression triggers the differentiation of cultured epidermal stem cells^17^. Deletion of the transmembrane protein collagen XVII (COL17), cytoplasmic integrin-linked kinase (ILK) or kindlin, which mediate the linkage between the epidermis and BM, resulted in defects in epidermal tissue regulation^18, 19, 20, 21, 22^. In addition, previous analyses showed differences in the expression of ECM genes between distinct subpopulations of epidermal stem cells^12, 15, 23^. These different ECM components may serve to anchor specific stem cells in the niche, and may also play a role in communication between epidermal stem cells and adjacent dermal cell populations. Indeed, BM proteins derived from certain epidermal stem cell populations provide a niche for the development and functions of arrector pili muscles and mechanosensory nerve complexes^12, 15^. Thus, it is likely that both the epidermal and the dermal surfaces of the BM are molecularly and structurally tailored to mediate distinct inter-tissue interactions.

Here, we systematically identified the cellular origins, molecular identities and tissue distribution patterns of ECMs in the mouse HF at high spatial resolution in a semi-quantitative manner. Our study provides the first comprehensive overview of the ECM landscape within the adult HF and highlights how BM composition and structure are exquisitely tailored for individual inter-tissue interactions. Our study further revealed a remarkable degree of molecular complexity and spatial specialization of BMs in the HG–DP interface, which is involved in the structural and functional integrity of the HG–DP inter-tissue interactions.

## Results

### Global ECM gene expression profiling in adult mouse hair follicles

Deeper sequencing is required to obtain comprehensive genome-wide ECM gene expression profiles, including that of low-abundance genes. Thus, we pooled different epidermal and dermal cell populations from adult telogen dorsal skin using cell sorting. We used our recently established epidermal cell isolation protocol to purify basal epidermal stem/progenitor cells (integrin α6^+^) resident in the lower isthmus (*Lgr6*^+^), upper bulge (*Gli1*^+^), mid-bulge (CD34^+^), HG (*Cdh3*^+^) and unfractionated basal epidermal stem/progenitor cells (basal; mostly from the interfollicular epidermis) by utilizing *Lgr6-GFP-ires-CreERT2*, *Gli1-eGFP, Cdh3-eGFP* and wild-type mice (Fig. 1a–e, Supplementary Fig. 1a–e)^12^. Two dermal cell populations, DP cells (*Lef1^+^/Pdgfra^+^*) and pan-dermal fibroblasts (*Pdgfra*^+^), were also isolated from *Lef1-eGFP* and *Pdgfra-H2B-eGFP* mice (Fig. 1f, g, Supplementary Fig. 1f–l). The purity of the sorted cells was verified by qRT-PCR on the HF region-specific genes^12^ (Fig. 1h). Then, RNA-seq of each isolated population was performed. Spearman’s rank correlation coefficient analysis and principal component analysis (PCA) with all of the expressed genes showed that all biological replicates clustered together and were significantly different from other samples (Supplementary Fig. 1m, n).

To investigate global differences in ECM gene expression levels in each cell population, we examined the FPKM values of all 281 annotated ECM genes, called the ‘matrisome’ (see Methods), in distinct cell populations^6^. Although the median FPKM of all genes in each cell population were around 1, those of the matrisome tended to increase from basal (median FPKM ~0.115) to HG populations (median FPKM ~0.958) (Fig. 1i). The median FPKM values of the matrisome in DP cells and dermal fibroblasts were ~2.14 and ~4.88, respectively, reflecting their roles in the production and maintenance of abundant interstitial ECM proteins. The diversity of the expressed ECM genes was also increased as cells were localized deeper in the skin (Fig. 1j). The number of expressed BM genes was constant across the examined populations, while that of interstitial matrix genes was increased in the deep portion of the HF.

The global ECM gene expression correlations among these cell populations were examined by hierarchical clustering. When all expressed genes were used, the epidermal and dermal populations were clearly separated (Fig. 1k). Intriguingly, however, when matrisome genes were used, HG was clustered with DP, but not with other epidermal cell populations, demonstrating that the ECM expression profile of HG cells resembles that of DP rather than those of other epidermal populations. This result also revealed that the ECM profile of DP resembles that of HG cells rather than that of pan-dermal fibroblasts.

Taken together, these results indicate that the matrisome of epidermal basal stem/progenitor subpopulations gains more complexity for epidermal cells located deeper in the skin. Given that the HG matrisome shows a particular resemblance to that of DP, ECM expression profiles of epidermal stem/progenitor compartments may be coupled with that of the adjacent tissues to cooperatively establish extracellular microenvironments, or niches, for local inter-tissue interactions.

### Cellular origin of BM and interstitial ECM molecules

It has been generally believed that the epidermal cells are the major source of the epidermal BM. However, no comprehensive understanding of the cellular origin of BM components has been obtained. Here, we systematically and quantitatively examined the expression patterns of 67 BM genes and 214 interstitial matrix genes in the isolated cell populations (Fig. 2, Supplementary Table 1). Eighteen BM genes (*e.g. Lama1, Lama3, Lama5, Col4a3, Col4a4, Col4a5, Col4a6, Col17a1, Egfl6, Fras1, Frem2, Npnt*) were exclusively or predominantly expressed in the epidermal stem/progenitor populations (Fig. 2, Supplementary Table 2). These genes can be classified into two categories: those encoding molecules functioning toward the epidermis (*Lama3, Lama5, Col17a1* – key molecules for keratinocyte adhesion)^24^ and toward the dermis (*Egfl6, Fras1, Frem2, Npnt* – key molecules for epidermal–dermal interactions)^12, 15, 25^. In contrast, 23 BM genes (*e.g. Lama2, Lama4, Lamc3, Col4a1, Col4a2, Col6, Col15a1, Nid1, 2, Ntn1–5*) were exclusively or predominantly expressed in the dermal fibroblast populations. This group of genes contains core BM genes, *Col4a1, Col4a2, Nid1* and *Nid2*. Other notable ECM genes were *Lama2*, *Lama4* and *Col6* isoforms, which mainly function for mesenchymal cells, such as nerves, muscles and blood vessels^24^. Eighteen BM genes (*e.g. Lamb2, Lamc1, Col7a1, Col18a1, Hspg2, Agrn, Sparc, Tgfbi, Tnc*) were expressed by both compartments. Together, our data indicate that major BM molecules for keratinocyte adhesion are provided by basal keratinocytes themselves, and that the dermal fibroblasts are another major source of BM molecules.

**Fig. 2.**
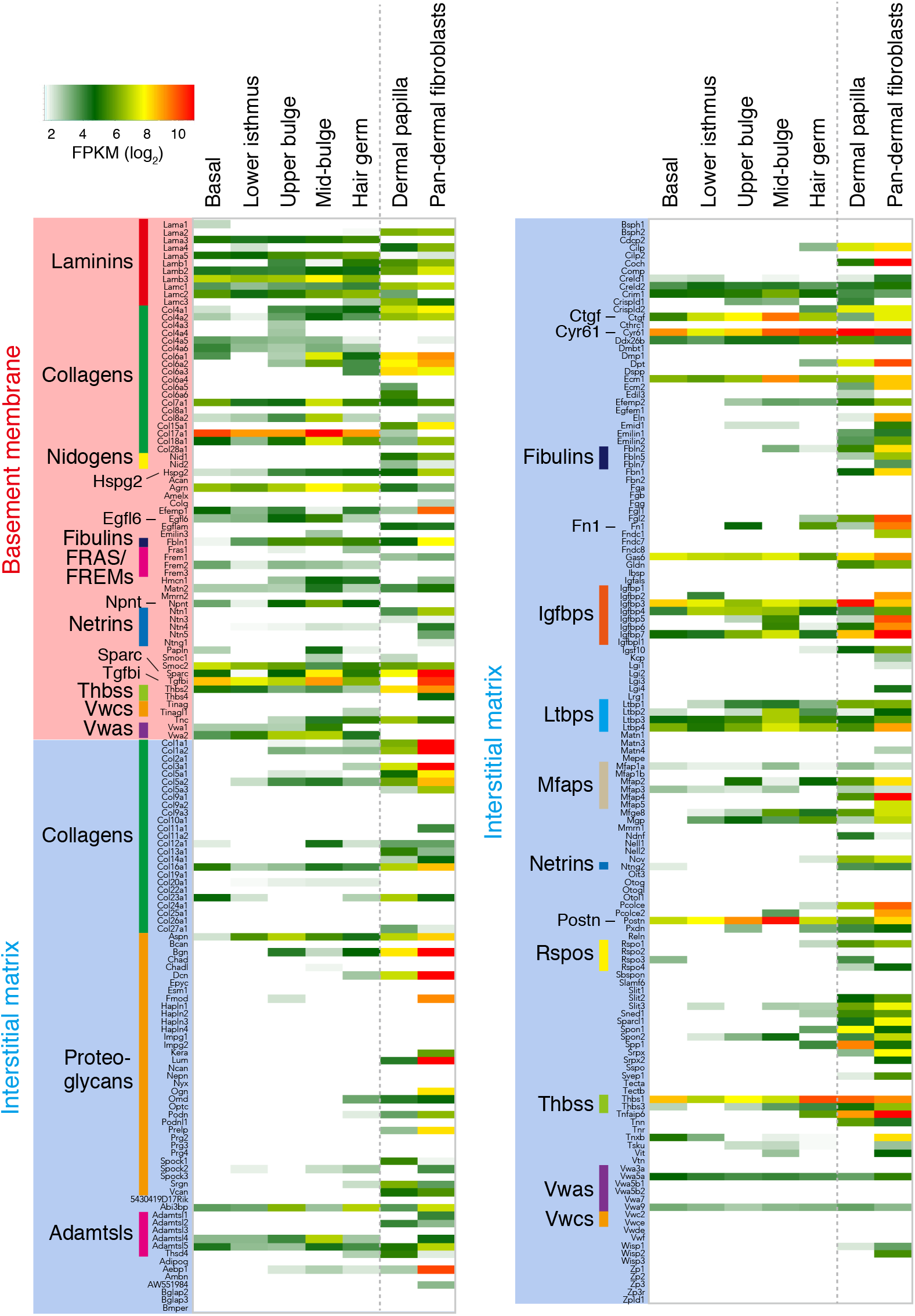
Overview of the matrisome gene expression. Heatmap representation of the expression levels of all matrisome genes. Averaged FPKM values (log_2_ converted; three or four biological replicates) are indicated according to a heat gradient as shown at the top left corner. Matrisome genes are first categorized into BM components (red zone) and interstitial matrix components (blue zone). Then, they are subdivided according to the ECM families to which they belong. Epidermal and dermal compartments are separated by a dashed line.

Interstitial matrix genes were expressed mainly by dermal cell populations (Fig. 2). Dermal fibroblasts expressed massive amounts of major interstitial structural ECM molecules such as *Col1a1, Col3a1, Col5a1, Col16a1, Bgn, Dcn, Lum, Fn1*, *Igfbp, Postn, Spp1* and *Thbs1*. Interstitial proteoglycans were almost exclusively expressed in dermal fibroblasts. Notably, however, a substantial number of interstitial matrix genes were also expressed by distinct epidermal cells (*e.g. Aspn, Abi3bp, Adamtsl4, Ctgf, Cyr61, Ecm1, Gas6, Igfbps, Ltbps, Postn, Thbss). Postn* and *Aspn* were highly expressed in the mid-bulge and upper bulge epidermal stem cells, respectively, and their protein products have been shown to be accumulated in the sub-BM zone of the mid-bulge and upper bulge regions^12, 15^. Thus, epidermis-derived interstitial matrix molecules could determine the region-specific distribution of interstitial matrix molecules at sub-BM zones, despite them being highly and broadly expressed in the dermis.

### ECM genes commonly or differentially expressed among different epidermal stem/progenitor cell populations

To identify ECM genes commonly or differentially expressed among different epidermal stem/progenitor cell populations, we categorized ECM genes based on their cell-type-dependent expression patterns. We identified cell populations that express certain ECM genes significantly more than other cell populations based on the following criteria: a gene expression level in a cell population i) of FPKM value ≥3 and ii) more than 40% of the FPKM value of the most highly expressed cell population. Categorized genes were depicted using a Venn diagram (Fig. 3a, Supplementary Table 3). We identified a commonly expressed epidermal stem/progenitor ECM gene group at the middle of the Venn diagram (group 1 in Fig. 3a). This gene group included *Lama3, Lama5, Lamb2* and *Lamc1*, which are components of laminin-332 and laminin-521, the laminin isoforms important for basal keratinocyte adhesion to the BM^26, 27^. *Crim1* is a transmembrane ECM gene and its defect results in embryonic skin blebbing and syndactyly^28, 29^. For these common ECM genes, ECM adhesion- and hemidesmosome assembly-related Gene Ontology terms were overrepresented (Fig. 3b).

**Fig. 3.**
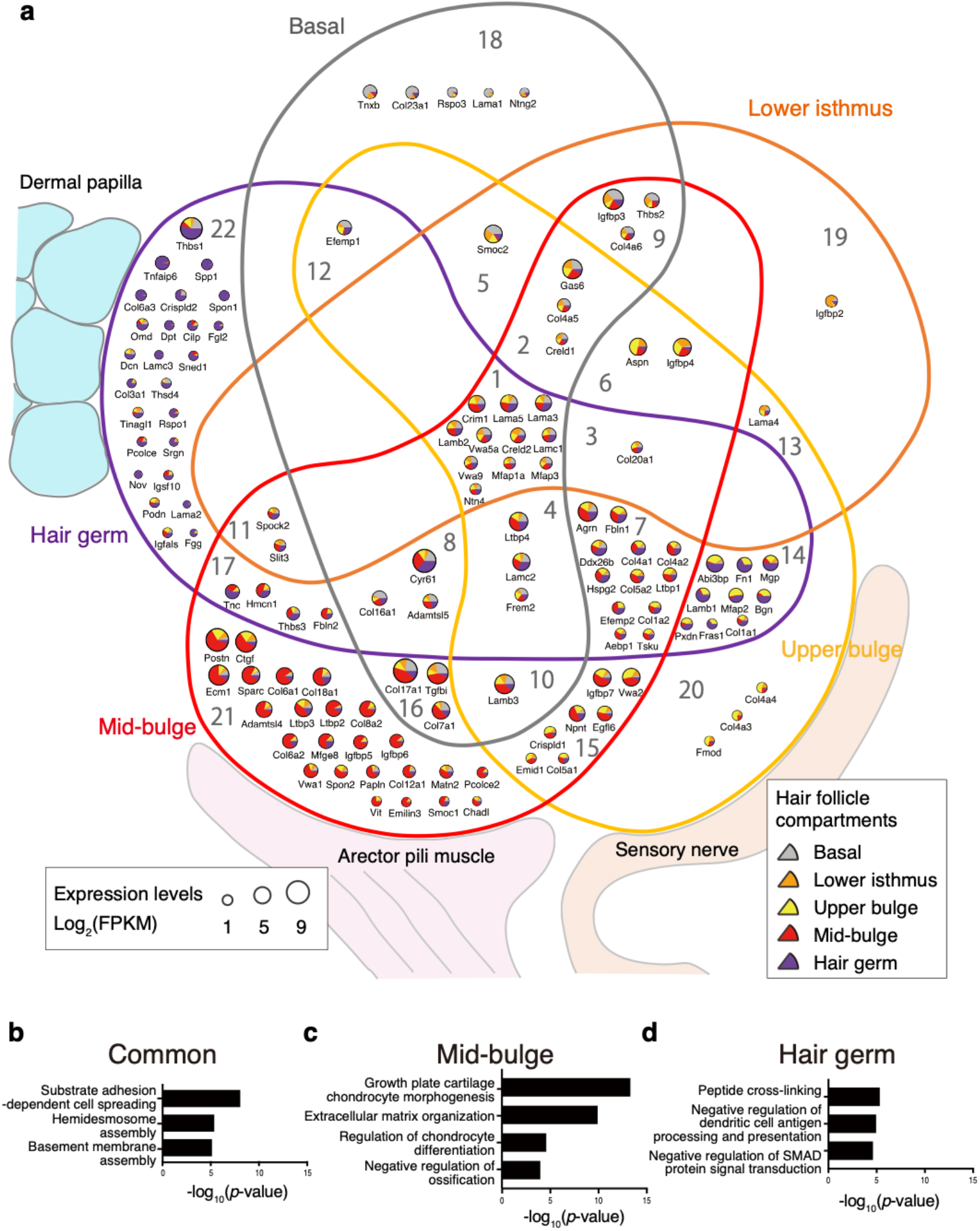
Classification of ECM genes based on their expression patterns in different epidermal cell compartments. **a** Venn diagram representation of the classification of ECM genes based on their expression in the epidermal cell compartments. Expression levels and ratios of each ECM gene are illustrated as a pie chart. The size and colours of the pie chart indicate the gene expression levels (lower left) and expressed compartments (lower right), respectively. To determine the presence or absence of regional expression, first, the maximum FPKM value of each gene among five epidermal compartments was obtained and expression levels relative to the maximum value were calculated for FPKM values in the other compartments. The threshold for adequate gene expression was set at more than 40% of the maximum FPKM and FPKM=3. Then, ECM genes were mapped in a Venn diagram based on their binarized expression data. Group numbers of ECM genes are marked in each category (see Supplementary Table 3). **b-d** Enriched GO terms analysed using common (b) or region-specific ECM genes (c, d).

Region-specific ECM genes were sorted at the periphery of the Venn diagram. Our analysis identified 24 mid-bulge-enriched ECM genes (group 21). Gene Ontology analysis of these mid-bulge ECM genes revealed that they were associated with cartilage or chondrocyte morphogenesis (Fig. 3c), suggesting the production of an ECM niche for the interaction with muscles. Indeed, one of the mid-/upper bulge-derived ECM proteins (group 15), nephronectin (*Npnt*), has been shown to play critical roles in arrector pili muscle development^15^. *Col17a1*, which is an important transmembrane ECM component for HF and interfollicular epidermal stem cell maintenance, was also identified as a mid-bulge-/basal-specific ECM gene (group 16)^18, 21, 22^. Sorting of these functionally important ECM genes into corresponding ECM gene groups demonstrates the reliability of the analysis.

Our analysis also identified 25 unique ECM genes highly enriched in HG (group 22). The SMAD signalling-related Gene Ontology term was overrepresented for these ECM genes (Fig. 3d). For example, *Thbs1* (thrombospondin-1), *Tnfaip6* (TSG-6) and *Spon1* (spondin-1/F-spondin) are involved in many morphological processes through TGF-β family signal regulation^30, 31, 32^. Consistent with these findings, dermal-derived TGF-β2 is critical for the activation of HG cells during the hair cycle^33^. Another key signalling pathway for HG–DP interactions is the Wnt/β-catenin signalling pathway^34^. *Rspo1* (R-spondin), an agonist of Wnt/β-catenin signalling^35^, was identified as an HG-specific ECM component. These results demonstrate that HG stem/progenitor cells express ECM genes involved in morphogen/growth factor regulation.

The expression patterns of ECM receptor genes were also examined. Laminin binding receptors, *Itga3, Itga6, Itgb4, Itgb1, Dag1* and *Bcam*, and their associated tetraspanins were highly and broadly expressed in epidermal stem/progenitor cells (Supplementary Fig. 2). Integrins for interstitial ECM molecules, such as *Itga5* (fibronectin receptor), *Itga1* and *Itga2* (collagen receptors), were enriched in HG cells, consistent with their high expression of interstitial matrix genes (Fig. 1j). In comparison to ECM genes, ECM receptor genes show broader expression patterns in the epidermis.

Taken together, these results indicate that all epidermal stem/progenitor cells commonly express ECM genes that are involved in epidermal–BM adhesion, while each epidermal stem/progenitor cell expresses region-specific ECM genes that play important roles in regional epidermal–dermal interactions.

### ECM genes commonly or differentially expressed among different fibroblast populations

ECM genes expressed in fibroblast populations were categorized in the same manner as in the epidermis (Fig. 4a, Supplementary Table 3). Although DP cells are a subpopulation of dermal fibroblasts, they shut down the expression of many major interstitial ECM genes, including *Dcn, Col1a1, Col1a2, Sparc, Col3a1* and *Lum*. Instead, they expressed ECM genes highly expressed in HG epidermal cells (*i.e. Spp1, Spon1, Lamc3, Srgn, Thsd4*) (Fig. 4b). Spondin family genes, including *Spon1, Rspo2* and *Rspo3*, were also upregulated, suggesting their roles in localized Wnt signal regulation.

**Fig. 4.**
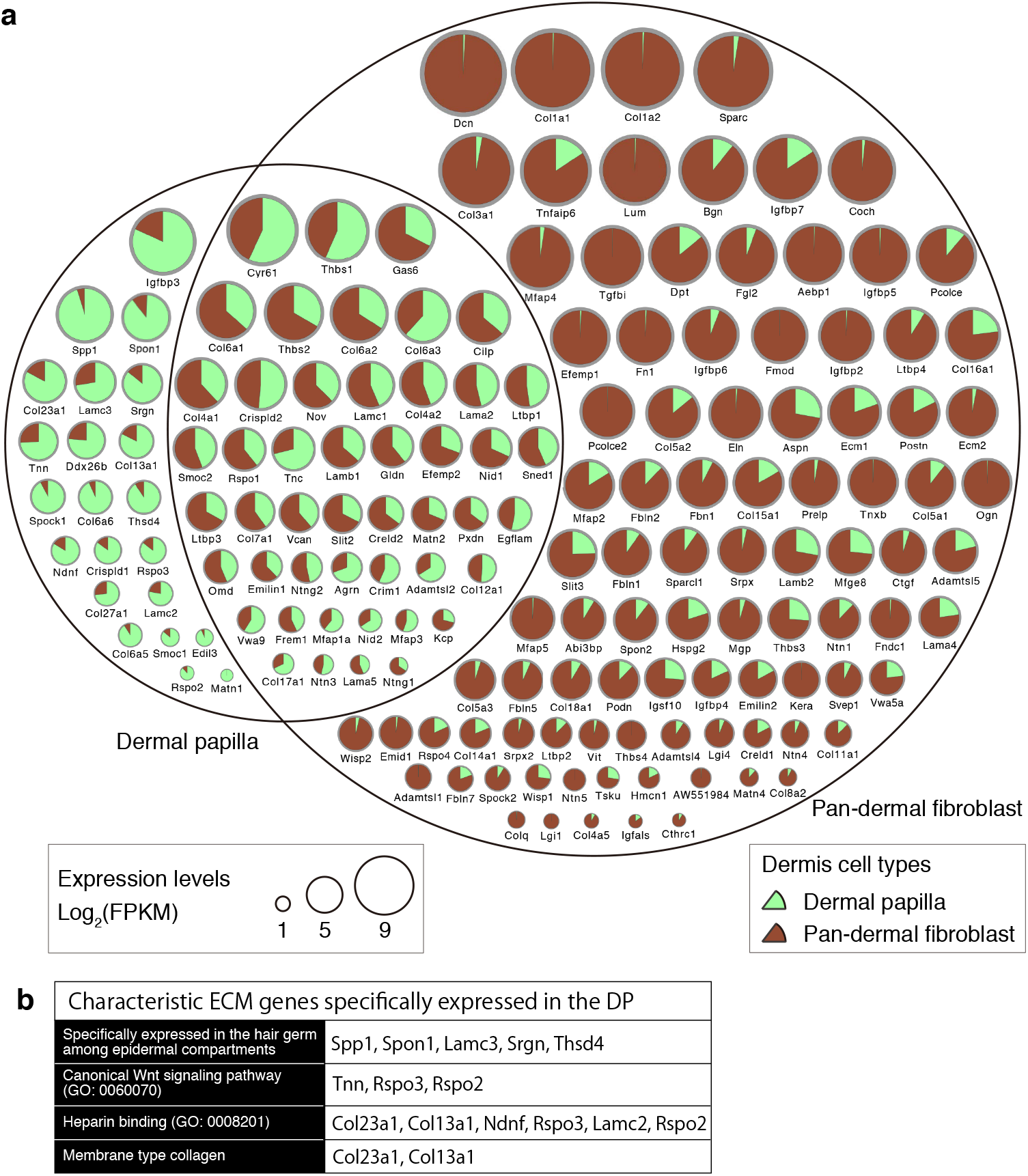
Classification of ECM genes based on their expression patterns in different dermal compartments. **a** Venn diagram representation of the classification of ECM genes based on their expression in the dermal cell compartments. The pie chart visualization and binarization methods are the same as in Figure 3a. **b** Characteristic ECM genes specifically expressed in the DP. Of the DP-specific ECM genes in the dermis, five are also found in the HG-specific ECM gene group in the epidermis. Many other DP-specific ECM genes are categorized in biological process GO terms related to growth factor regulation (Canonical Wnt signalling pathway and Heparin binding).

### ECM protein tissue atlas of mouse hair follicles

We further examined the tissue localization of commonly or regionally expressed epidermal and dermal ECM proteins by immunostaining and generated an ECM protein tissue atlas of mouse HFs. We used antibodies against 66 ECM proteins (tested 90 antibodies) and determined their immunostaining conditions with rigorous validation for specificity. Among them, 41 antibodies showed ECM-like extracellular deposition patterns (Supplementary Table 4). Immunostaining patterns of representative ECM proteins for each HF compartment are shown in Figure 5a and those of all 39 ECM components are shown in Supplementary Figure 3. Protein deposition levels of 34 ECM proteins were quantified and represented as a heatmap (Fig. 5b) and radar charts (Fig. 5c). These data clearly demonstrated that most ECM proteins detected showed tissue deposition patterns consistent with their gene expression patterns. For example, commonly expressed ECM components, such as laminin-332 (*Lama3*), laminin α5 (*Lama5*) and laminin β2 (*Lamb2*), were broadly detected in the BM zone of the interfollicular epidermis and HF (Fig. 5a–c). In contrast, ECM proteins specifically expressed in upper bulge, mid-bulge, HG and DP were preferentially deposited in their own cell compartments (Fig 5a, b, d–g), indicating that most ECM proteins are locally synthesized and deposited into matrices.

**Fig. 5.**
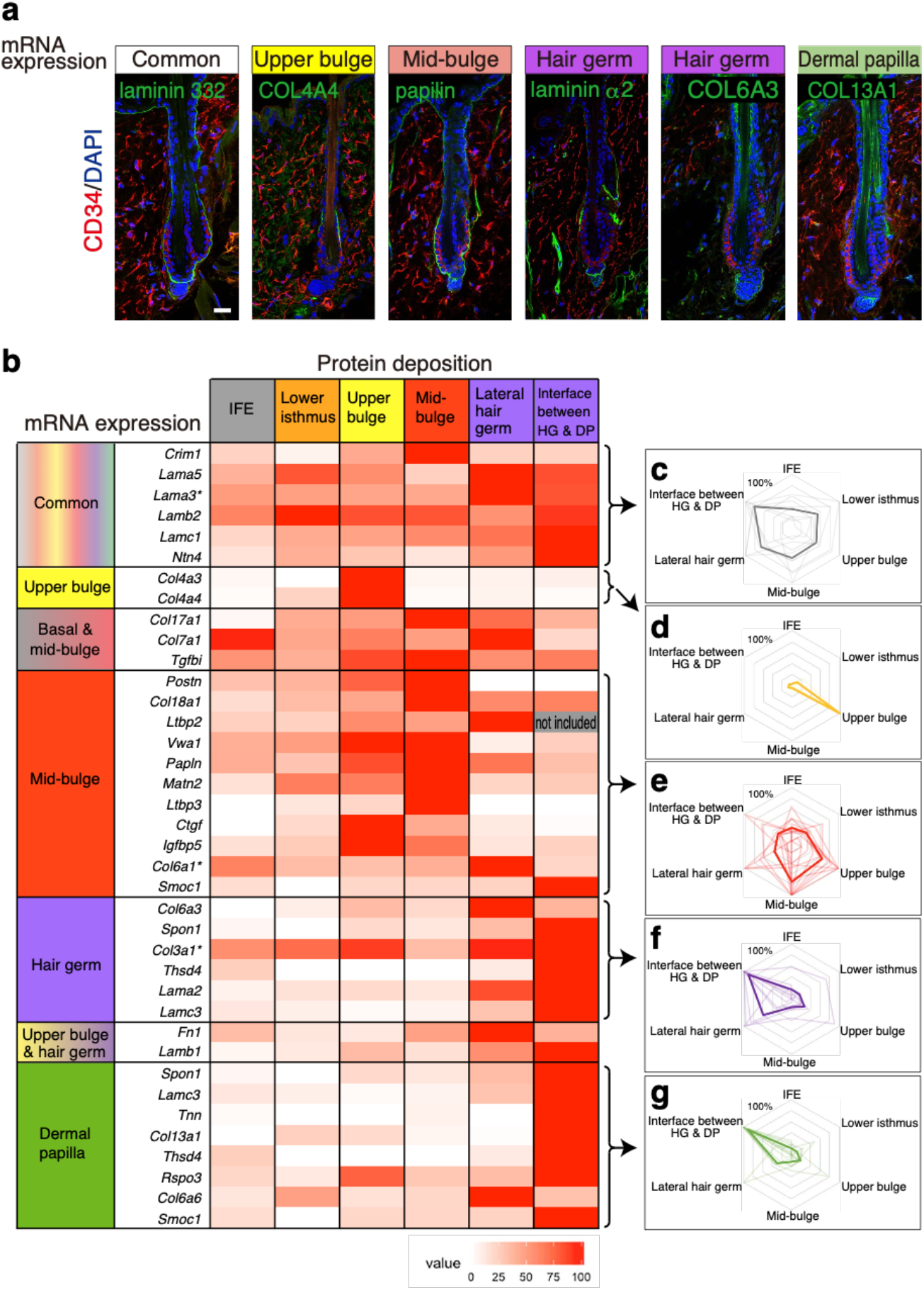
Specialized ECM niches and their cellular origins in the adult telogen hair follicle. **a** Representative tissue localization patterns of ECM proteins expressed as common or region-specific ECM genes. Immunofluorescence detection of each target ECM protein (green) and CD34 (red) in dorsal telogen HFs is shown with DAP1 counterstaining (blue). Scale bar, 20 μm. **b** Heatmap displaying the quantified deposition levels of ECM proteins. Using immunofluorescent histochemical staining data (Fig. 5a, Supplementary Fig. 3), signal intensities of ECM proteins deposited at the divided BM regions were measured as percentage values and depicted in a heatmap. Skin BM was regionalized into six areas as shown at the top of the heatmap. The major cellular origins of each ECM molecule are indicated at the left panel of the heatmap. Asterisks on the gene symbol indicate the use of an antibody that does not distinguish a subunit composing ECM protein complexes. IFE: interfollicular epidermis. **c–g** Radar chart analysis of protein tissue distributions of region-specific ECM genes. Each radar chart consists of six BM regions. Quantified deposition levels of ECM proteins in (b) are plotted and represented as thin lines and their average patterns are depicted by a bold line [common in (c), upper bulge in (d), mid-bulge in (e), HG in (f) and DP in (g)].

Each column of the heatmap defines the ECM niche of each cellular compartment and clarifies the molecular differences (Fig. 5b and Supplementary Fig. 4). The epidermal BM showed a clear boundary of ECM deposition, like a patchwork, even though the BM is a continuous sheet-like structure. Many region-specific ECM proteins showed consistent deposition borders with those of CD34 bulge marker expression, suggesting that ECM environments are specialized along epidermal cellular compartments. We also found that the ECM composition clearly differed between HG lateral BM and HG–DP interface BM, a finding that is explored further below.

### Identification of BM micro-niches along epithelial–mesenchymal interfaces

We next probed the diversity within the BM niches in the HG–DP unit, which governs HF morphogenesis and regeneration, with the ECM protein tissue atlas. The first notable feature that we found was the lack of reticular lamina components in the BM forming the interface between HG and DP. The reticular lamina components, COL6 (*Col6a1, α3, α6*) and COL7 (*Col7a1*), were absent or present at very low levels at the interface BM (Fig. 6a arrows). The hemidesmosome component COL17 (*Col17a1*) was also absent from the interface BM (Fig. 6a arrow). We thus postulated that the interface BM has altered BM and/or hemidesmosome structures. We then examined the expression patterns of the intracellular hemidesmosome protein plectin and found that it was also absent from the interface BM zone (Fig. 6b arrow). Ultrastructural examination revealed that the number of electron-dense hemidesmosome-like structures was reduced at the interface BM (Fig. 6c–f). Notably, the lamina densa structure at the interface BM showed protrusions toward the dermis (Fig. 6g arrows), while these protrusions were not observed in the lateral BM. These lamina densa protrusions preferentially originated from hemidesmosome-like structures (Fig. 6g arrowheads). It has been reported that the BM of the neuromuscular junction also lacks reticular lamina and extends protrusions from active zones to junctional folds of muscle fiber^36, 37^. Thus, our analysis identified close parallels in molecular composition and structure of the BM between the HG–DP interface and neuromuscular junction.

**Fig. 6.**
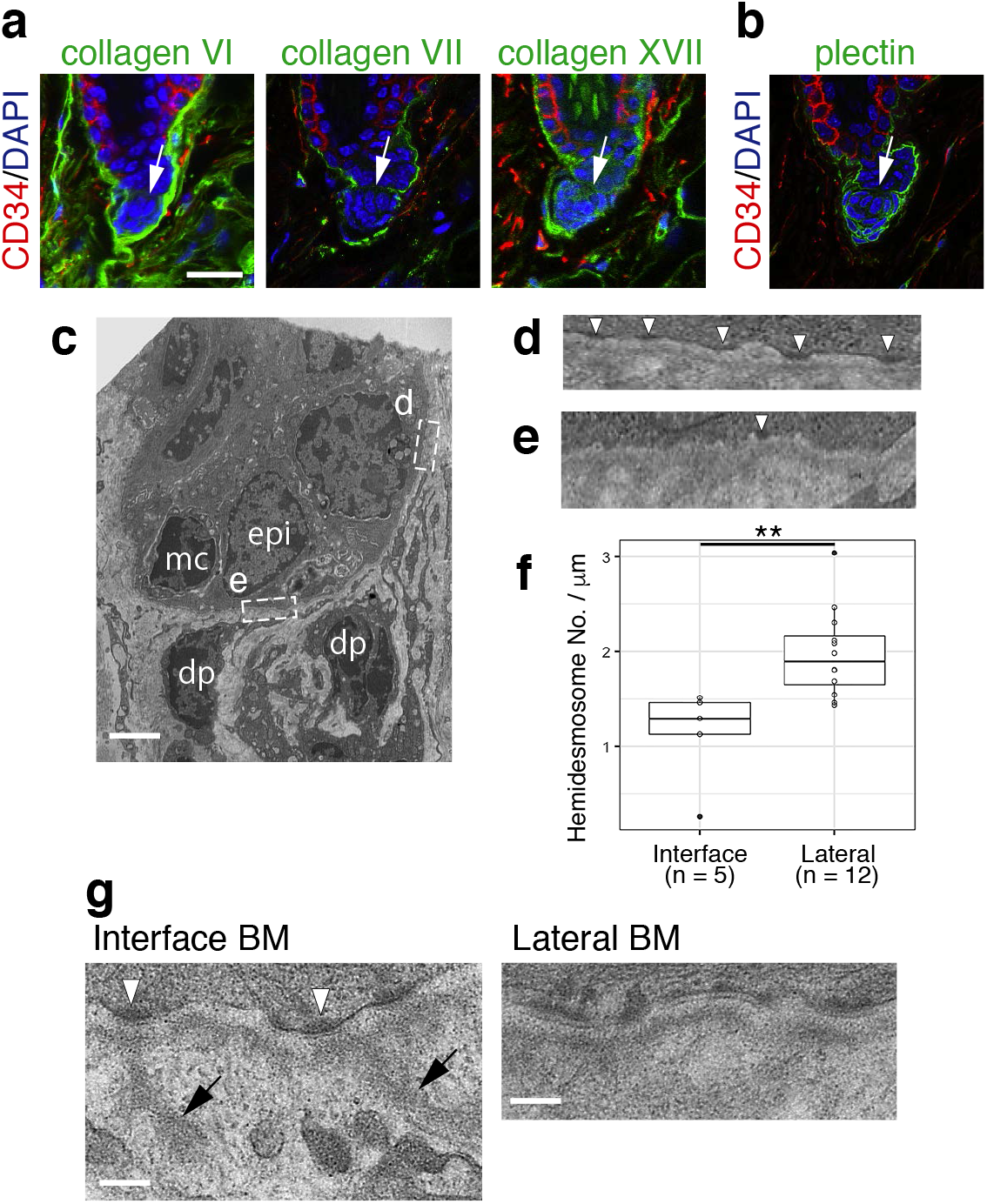
Absence of reticular lamina, fewer hemidesmosomes and extension of BM protrusions at the hair germ and dermal papilla interface. **a, b** Immunolocalizations of reticular lamina-related ECMs, COL6 and COL7, and hemidesmosomal components, COL17 (a) and plectin (b), in dorsal telogen HFs. These ECMs and plectin (green) were co-stained with CD34 (red) and DAPI (blue). White arrows indicate the interface between HG and DP. **c** Transmission electron microscopy (TEM) image of HG and DP region. epi, epidermal HG cell; mc, melanocyte; dp, DP cell. **d, e** Magnified images of HG–BM adhesion sites located at the lateral side (d) and interface side (e) of the HG region (c). Hemidesmosome structures are indicated by white arrowheads. **f** Box plot of the hemidesmosome densities of HG cells located at the lateral or interface sides of the HG region. ** *p* <0.01, Mann–Whitney U test (two-tailed). **g** BM protrusions observed at the interface BM. BM protrusions extending into the interstitial space are marked with arrows. Hemidesmosome structures are indicated by white arrowheads. Scale bars: 20 μm (a), 2 μm (c), 500 nm (g).

We further identified two additional specialized BM structures in the HG–DP unit. Laminin α5 staining showed large protrusions from the interface BM into the centre of the DP where a nuclear signal was lacking (Fig. 7a). Another core BM molecule, perlecan, overlapped with laminin α5, indicating that these protrusions are continuous extensions of the BM. Three-dimensional image analysis showed that this ECM structure resembles a hook that fastens DP to the HG (Fig. 7b). Thus, we named this novel ECM structure the ‘hook BM’. The hook BM also contains other major BM molecules, including laminin α2, α4, β1, β2, γ1, γ3, nidogen-1, −2 and COL4, but not laminin-332, suggesting that the major laminin isoforms in the hook BM are laminin α2, α4 and α5 chain-containing laminins (Fig. 7c). Among them, γ3 chain-containing laminin isoforms are unable to bind to integrins^38^, suggesting their role in regulating integrin binding properties of hook and interface BMs. We also noticed a mesh-like deposition of perlecan within the DP (Fig. 7a). This mesh-like ECM structure extended to the interspace among DP cells and was directly connected to the interface and hook BMs. We named this ECM structure the ‘mesh BM’. The mesh BM also contains laminin α4, β1, γ1, nidogen-1, −2 and COL4, but not laminin α2, α5, β2, γ3 and laminin-332, suggesting that the major laminin isoform of the mesh BM is laminin-411. We also found that other BM molecules, such as netrin-4, LTBP3, adamtsl-6, tenascin-N and COL13A1, were located in the hook BM, while spondin-1, adamtsl-6, biglycan, tenascin-N and R-spondin-3 were localized in the mesh BM (Supplementary Fig. 5a). Both epidermal and dermal cell compartments contribute to producing these BM components (Fig. 2). Taking these findings together, the ECM protein tissue atlas revealed exquisite molecular and structural diversity of BM micro-niches at the HG–DP interface (Fig. 7d) and showed that the BM is the primary ECM niche for DP cells.

**Fig. 7.**
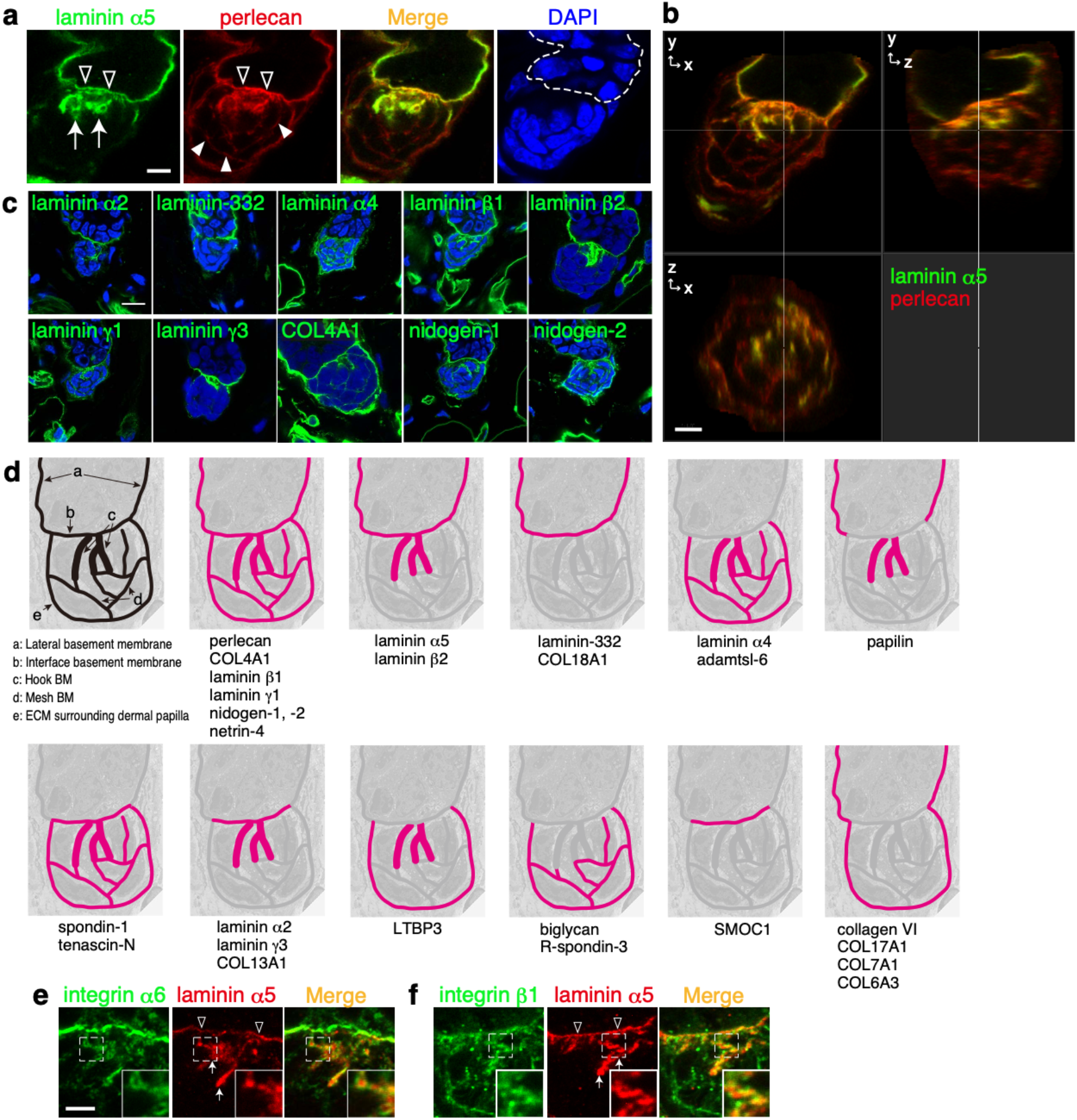
Identification of hook and mesh BMs in the hair germ–dermal papilla unit. **a** Immunofluorescence labelling of dorsal telogen HFs. Both laminin α5 (green) and perlecan (red) are detected in a hook-shaped BM (arrows) extending from the interface BM (open arrowheads). Perlecan also forms a mesh–like BM (filled arrowheads) within the DP. Dashed line indicates the epidermal-dermal boundary. **b** Three-dimensional observation of the hook and mesh BMs in (a). **c** Immunolocalizations of major BM proteins (green) in the hook and mesh BMs of dorsal telogen HFs. Nuclei were stained with DAPI (blue). **d** Graphical representation of the regional BM compositions in the HG–DP unit. Upper left-hand panel depicts the distinct BM structures. Other panels schematically summarize deposition patterns of BM components examined in a, c and Supplementary Fig. 3. **e, f** Close localization of dermal integrins with laminin α5-containing hook and interface BMs. Integrin α6 (e, green) or βl (f, green) was co-immunostained with laminin α5 (red). Insets are magnified views of the dotted squares. Arrows and open arrowheads indicate the hook BM and interface BM, respectively. Scale bars: 5 μm (a, b), 10 μm (c), 3 μm (e, f).

To identify potential ECM receptors for these BMs, we examined the expression levels of major ECM receptors in DP and pan-dermal fibroblasts. We found that DP cells highly expressed laminin-binding integrins (*Itga6* and *Itga3)^39^* and Wnt signal regulator glypicans (*Gpc1* and *Gpc2*)^40, 41^ (Supplementary Fig. 5b). Immunohistochemical detection of integrins showed that the integrins α3, α5, α6, α8, β1 and β4 were enriched at the interface BM region, α6, αv and β1 were located at the hook BM region, while integrins α5, α6, α9, αv and β1 were enriched at the mesh BM region (Supplementary Fig. 5c). Super-resolution microscopic images revealed that the integrins α6 and β1, which form an integrin α6β1 heterodimer that binds to laminin α5 chain-containing laminins^39^, tightly associated with laminin α5 in the interface and hook BMs, suggesting that integrin α6β1 on DP cells interacts with the interface and hook BMs (Fig. 7e, f).

To investigate the interactions between BMs and dermal cells, we performed electron microscopic analysis. Ultrastructural analysis visualized the close associations of BM protrusions and cellular protrusions of DP and dermal stem cells at the interface and hook BM regions (Supplementary Fig. 6a–e). In contrast, DP cells located away from the interface and hook BMs, where the mesh BM is located, exhibited relatively smooth and flat cell membrane structures (Supplementary Fig. 6f, g). Cell–cell interactions among DP and dermal stem cells were rarely observed; instead, the interface, hook and mesh BMs cohered these dermal cells. These results demonstrated that epidermal HG cells, DP cells and dermal stem cells are aggregated by tangling with a continuous BM structure, which have regionally specialized molecular compositions and structures. Taking the findings together, our analysis revealed a remarkable degree of molecular and structural complexity of the BM niches and a variety of cell–BM interactions at the HG–DP interface. The findings also indicate the existence of asymmetrically organized side-specific heterogeneity in BM composition and structure in this inter-tissue interface.

### Epidermal laminin α5 is required for the topological and functional integrity of the hair germ–dermal papilla unit

Laminin α5 appeared to be a major cell adhesion ligand for both HG and DP cells, and has also been reported to be involved in HF morphogenesis^27, 42^, therefore, we examined the effects of the deletion of the *lama5* gene at this interface. To investigate the cellular origin of laminin α5 at the interface, *Lama5* floxed mice were crossed with Keratin 5-Cre mice, which specifically express Cre in the basal epidermal compartment. In the mutant, immunoreactivity of laminin α5 disappeared from the interface and hook BMs, demonstrating that the laminin α5 in these BMs is derived from the epidermis (Fig. 8a). This result is consistent with our transcriptome data showing that *Lama5* is of epidermal origin (Fig. 2, Supplementary Table 2). The deletion of *Lama5* resulted in the failure in forming the bell-shaped epidermal structure at the tip of the developing whisker HFs (Fig. 8b). Furthermore, mutant DP did not show a prolate spheroid structure, but retained a round shape. Thus, the geometry of the HG–DP interface was significantly altered in the mutant, indicating that laminin α5 is an important regulatory element of tissue topology and cellular arrangement of the HG–DP unit.

**Fig. 8.**
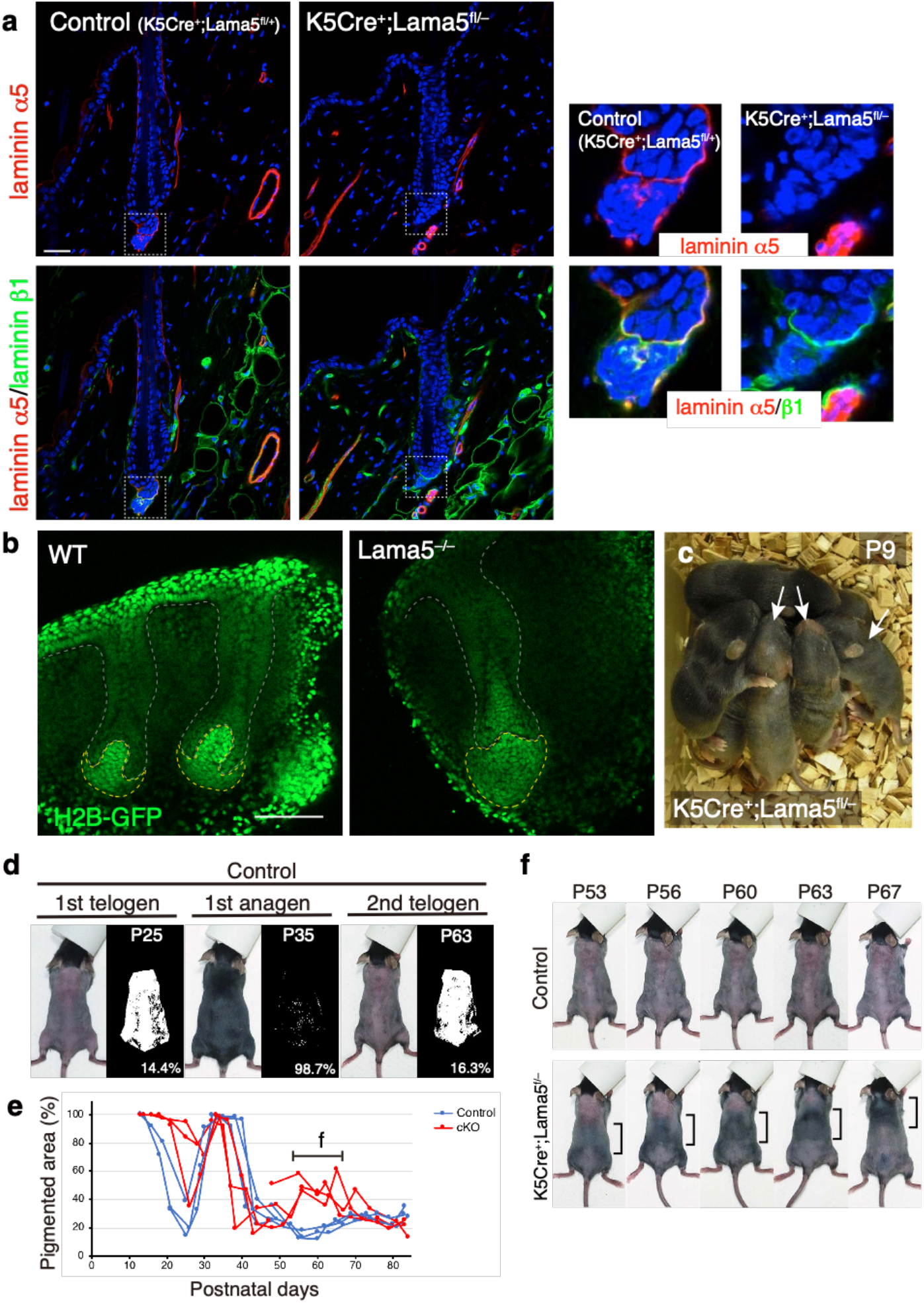
Laminin α5 is required for the topological and functional integrity of hair germ–dermal papilla unit. **a** Immunohistochemical examination of laminins α5 (*red*) and β1 (*green*) in the dorsal telogen HFs of control (K5Cre^+^;Lama5^fl/+^) and K5Cre^+^;Lama5^fl/−^ mice. Magnified images of the dotted squares are shown (*right panels*). **b** Fluorescent images of developing whisker HFs in E14.5 R26-H2B-GFP (WT) and R26-H2B-GFP;Lama5^-/-^ embryos. Grey dotted lines indicate the border between epidermis and dermis. Yellow dotted lines indicate the border of dermal papilla. **c** Gross appearance of newborn K5Cre^+^;Lama5^fl/−^ mice (*arrows*) with their control littermates. Hair growth is delayed. **d** Measurement of pigmented skin areas. Pigmented area, which is associated with the presence of anagen HFs, on the shaved dorsal skin was determined from binarized photos (right side of each panel) at various postnatal days (P). **e** Comparison of hair cycle patterns between control and mutant mice. Ratio of pigmented area represents the ratio of anagen HFs. Four litter pairs were examined. One pair of littermates was examined from P21 to P47 and another pair of littermates was examined from P48. A horizontal bar indicates the period for detailed hair cycle pattern analysis shown in f. **f** Precocious anagen entry and the formation of traveling hair regeneration wave in K5Cre^+^;Lama5^fl/−^ mice. Scale bars: 100 μm (b).

In accordance with the anatomical defects, hair growth was significantly reduced in the newborn mutant mice (Fig. 8c). We also investigated the continuum of hair cycle stages by observing hair cycle domain patterns on living mice^43^. In the mutants, the onset of the first catagen (regression phase between the growing anagen phase and resting telogen phase), was delayed, while HFs entered the next anagen at the same timing as the control mice without showing clear telogen transition (Fig. 8d, e). In the second telogen phase (~P45–80), the mutants showed precocious anagen entry and exhibited a hair regenerative wave traveling from the tail to the head in the dorsal skin (Fig. 8e, f), suggesting the misregulation of signalling events for cyclic HF regeneration. Taken together, these results indicate that epidermal-derived laminin α5 at the HG–DP interface plays a key role in the topological and functional integrity of the HG–DP unit.

## Discussion

One of the important issues regarding the ECM is how the ECM composition and structure are spatiotemporally specialized to orchestrate organ formation and function. However, the entire molecular landscape of the ECM composition and its pattern-forming processes in all organs remain largely unknown. Here, we systematically and thoroughly identified the cellular origins, molecular identities and tissue distribution patterns of ECMs in the mouse HF at high spatial resolution in a semi-quantitative manner. Our study provides the first comprehensive overview of the ECM landscape within the adult HF and highlights how ECM composition is regionally specialized for each cell type and distinct inter-tissue interactions.

### Mapping both transcriptome and protein tissue localization of the ECM with high spatial resolution

There are two major strategies to make regionally specialized ECMs: synthesis locally at a target cell/tissue or selective accumulation of ECMs from a distant source^3^. Our combinatorial systematic ECM mRNA and protein mapping approach revealed that, at least in the epidermal–dermal interface, most ECM molecules are synthesized locally and accumulated in adjacent ECMs, indicating that localized ECM expression is a major determinant of the ECM environment for both epidermal and dermal populations. This seems reasonable, but is also surprising because very different combinations of ECM proteins can be deposited locally together, despite the interaction and assembly of ECM molecules being considered to be regulated by specific molecular interactions^24^. This suggests the existence of molecular interaction networks within the locally expressed ECM molecules for their effective assembly and turnover. Our data also suggest that the ECM niche of each cell type can be well predicted from gene expression profiles of their own and neighbouring cells. This finding lays the foundation for predicting ECM niches of single cells within tissues by using large-scale single-cell gene expression datasets, such as the Human Cell Atlas^44^, Human Biomolecular Atlas Program (HuBMAP)^45^ and the mouse atlas *Tabula Muris^46^*.

Our analysis also identified ECM proteins that exhibited inconsistent tissue localization patterns between mRNA and protein. For example, *Crim1* was categorized as a commonly expressed epidermal ECM gene, but its protein product was enriched at the bulge. In addition, fibroblast-derived BM proteins such as laminin α2, α4 and COL4 also exhibited distinct localization patterns in the BM despite being highly expressed in a distant source, pan-dermal fibroblasts. These findings imply the existence of molecule-specific long-range ECM transport and assembly mechanisms, as reported previously in *Drosophila^47^*. Although it is unclear how their tissue localizations are regulated, ECM receptors and other interacting ECM proteins play roles in their tissue localization. Deeper and/or more diverse systematic molecular profiling and computational analysis of the expression and localization of ECM molecules and receptors will help us understand how distinct ECM niches are generated.

Our antibody-based ECM mapping revealed complex subcellular ECM distributions, leading to the identification of the hook and mesh BMs. These matrices form molecularly and structurally fine-tuned ECM niches at the HG–DP interface. This level of spatial mapping resolution cannot be achieved by other current proteome approaches such as mass spectrometry^48, 49^. Thus, merged antibody-based spatial ECM protein and single-cell mRNA expression profiles can precisely relate ECM compositions to the position of cells and molecules, thus providing distinct ECM niche information and a common anatomical reference for normal, aged and pathological tissue structures.

### The BM provides a tailored interface for distinct inter-tissue interactions

The BM can simultaneously function as both a tissue insulator and glue, keeping different cell populations in close vicinity of each other with a clear tissue boundary^3^. Our analysis clearly showed that the molecular composition and structure of the BM are specialized for distinct inter-tissue interactions. The mid-bulge BM is composed of ECM molecules related to chondrocyte morphogenesis for the interaction with arrector pili muscles^15^. Surprisingly, these chondrocyte-related ECM molecules are derived from epidermal bulge stem cells, but not from chondrocytes or related cell types, indicating that epidermal bulge stem cells actively participate in cooperative formation of the niche for epidermal-muscle inter-tissue interactions. To this end, bulge stem cells need to activate a transcriptional network for chondrocyte ECM expression. Indeed, *Sox9* and *Scx*, master transcription factors for chondrocytes and tenocytes, are highly expressed in the bulge stem cells, potentially contributing to establish a BM niche for epidermal– muscle interactions^15, 50, 51^. In contrast, HG cells express a very different set of ECM genes, including those related to growth factor signalling, such as SMAD/TGF-β signalling and Wnt signalling. These signalling pathways are critical regulators of HG– DP interactions and HF morphogenesis and regeneration^16^. This marked difference in ECM expression patterns in the adjoining epidermal compartments reflects their different counterpart tissues for inter-tissue interactions. Thus, one of the major reasons for the heterogeneity and compartmentalization of epidermal basal stem/progenitor cells and their BM composition is to establish distinct inter-tissue interfaces between the epidermis and dermis^10^.

The absence of reticular lamina may allow intimate inter-tissue interactions by providing laminin–integrin interactions at both epidermal and dermal sides of the BM. We found that the BM between HG and DP lacks reticular lamina components, such as COL6 and COL7. This BM extends many protrusions from epidermal hemidesmosome-like structures toward the DP and dermal stem cells and forms laminin–integrin complexes at the dermal side. An analogous BM structure can be observed in the neuromuscular junction, where the reticular lamina is excluded and the BM extends protrusions from synaptic active zones toward junctional folds that invaginate the postsynaptic muscle membrane^36^. Laminin–integrin interactions can be observed on both nerve and muscle sides of the BM and play critical roles in the formation and function of neuromuscular junctions^52, 53^. Moreover, COL7 is absent from lung alveoli, blood vessels and kidneys, where different tissues are tightly integrated via the BM that places laminins at both its sides^1, 2, 54, 55, 56^. Thus, it is likely that the absence of the reticular lamina enables the placement of laminins on both sides of the BM.

### Existence of BM micro-niches in the hair germ–dermal papilla unit

There are numerous reports on the identification of signalling molecules involved in HG–DP interactions. These interactions would be governed by the spatial organization of heterogeneous HG and DP cells and their specialized micro-niches^57^. However, only limited information is available on the molecular identities of the extracellular space in the HG–DP unit. Our study revealed that the molecular composition of BM niches in the HG–DP unit is exquisitely tailored at the cellular level, likely to allow for coordinated multi-lineage interactions. HG–DP interactions have the following key features: i) DP cells form a packed cluster despite them being scatter-prone fibroblasts, ii) DP cells constantly attach to the HG region despite HFs undergoing dynamic tissue regeneration and iii) HG and DP cells actively exchange signals via soluble factors such as Wnts, BMPs, FGFs and TGF-βs^16^. The mesh BM could help provide feature i) because the cell–cell interactions of DP cells were limited; instead, the mesh BM cohered laminin-receptor-expressing DP cells. The interface, mesh and hook BM complex can potentially underpin features ii) and iii) because these BMs were physically connected and preferentially composed of different adhesion and soluble signalling-related ECM molecules. In fact, the deletion of an epidermal-derived interface and hook BM molecule laminin α5 resulted in disruption of the topological and functional integrity of the HG–DP unit during HF development and regeneration. Laminin α5 has been reported to be involved in many morphological processes via regulating integrin-mediated cell adhesion and growth factor-mediated signalling events^58^. In the skin, laminin α5 regulates keratinocyte adhesion, proliferation and migration, and is also suggested to be involved in growth factor signalling in the HFs^42, 59, 60, 61^. Thus, laminin α5 could function as a direct adhesion target for both HG and DP cells and could also regulate the tissue distribution and activity of growth factors.

Inter-tissue interactions are essential for the development, regeneration and functions of most organs. They have their own tailored BMs as structural and functional interfaces of inter-tissue interactions. Thus, future work should further characterize the molecular and structural properties of distinct BMs and their dynamics in inter-tissue interactions and reveal their significance in the coordination of multi-lineage interactions. This work provides a paradigm for understanding the role of BM heterogeneity in mediating inter-tissue interactions in multicellularity.

## Author Contributions

K.T. designed and carried out experiments, analysed data and wrote the manuscript. H.M. designed and carried out experiments and analysed data. A.N. provided technical support. R.M. examined phenotypes of whisker follicles of *Lama5^-/-^* mice. J.H.M. provided *Lama5* floxed and knockout mice. K.S. provided antibodies against vwa1, papilin, TGFBI, adamtsl-6, abi3bp, laminin β2 and γ3. H.F. conceived the project, designed and supervised experiments, analysed data and wrote the manuscript.

## Acknowledgements

We thank Shigehiro Kuraku, Chiharu Tanegashima, Sean D. Keeley, Yuichiro Hara and Osamu Nishimura of the Laboratory for Phyloinformatics, RIKEN, for help with the RNA-seq and bioinformatics; Yasuko Tomono (Shigei Medical Research Institute) for antibodies against type IV collagens; Yoshiaki Hirako (Nagoya University) for antibody against type XVII collagen; Phillipe Soriano (Mount Sinai NY) for *Pdgfra-H2B-eGFP* mice (Pdgfra^tm11(EGFP)Sor^); Jose Jorcano (CIEMAT-CIBERER, Madrid, Spain) for K5-Cre mice; Takaya Abe (RIKEN BDR) for R26-H2B-EGFP mice; and RIKEN Kobe light microscopy and animal facilities for technical assistance. We also thank members of the Fujiwara laboratory for valuable reagents and discussions. This work was funded by RIKEN intramural funding, RIKEN ‘Epigenome and Disease’, JSPS KAKENHI (25122720, 26670533), JST CREST (JPMJCR1926), Uehara Memorial Foundation, Takeda Science Foundation and Cosmetology Research Foundation (all to H.F.). H.M. was supported by the RIKEN Junior Research Associate (JRA) program.

## Methods

### Mice

*Lgr6-GFP-ires-CreERT2, Gli1-eGFP, Cdh3-eGFP, Lama5* floxed, *Lama5* knockout mice have been described previously^12, 62^. *Lef1-eGFP* mice [STOCK Tg(Lef1-EGFP)IN75Gsat/Mmucd] were obtained from MMRRC. *Pdgfra-H2B-eGFP* mice (Pdgfra^tm11(EGFP)Sor^)^63^, K5-Cre mice^64^ and R26-H2B-EGFP mice (CDB0238K)^65^ were kindly provided by Dr Phillipe Soriano (Mount Sinai NY), Dr Jose Jorcano (CIEMAT-CIBERER, Madrid, Spain) and Dr Takaya Abe (RIKEN BDR), respectively. Mouse lines used for transcriptome analysis were backcrossed with C57BL/6N mice more than four times. R26-H2B-EGFP/*Lama5^-/-^* mice were crossed with C57BL/6 albino mice several times to avoid possible imaging interference from melanin deposition. For cell sorting and immunohistochemical analysis, wild-type C57BL/6N mice were used.

### FACS

The epidermal cell isolation procedure was as described previously^12^. DP cells and pan-dermal fibroblasts were isolated from the dorsal skin of 8-week-old female mice. The epidermal tissue was removed from the dermal tissue by scraping after trypsinization. The remaining dermal tissue was minced and treated with 2 mg/ml collagenase type I (Gibco, MD, USA) at 37°C for 2 h with gentle mixing. Single-cell suspension was obtained by repeated gentle pipetting and passed through a 40 μm cell strainer (Falcon, NC, USA). To eliminate haematopoietic and endothelial cells (lineage-positive cells; Lin^+^), the cell suspension was stained with PE-Cy7-conjugated antibodies for CD45 (eBioscience, CA, USA, 30-F11), TER-119 (eBioscience, TER119) and CD31 (eBioscience, 390). To further distinguish the target cell populations from the remaining epidermal cells, the expression of CD49f (integrin α6) was examined using a PE-conjugated antibody (eBioscience, GoH3). For the further analysis of dermal cell populations, the cell suspension was also stained with CD34-eFluor660 antibody (eBioscience, RAM34). Cell isolation procedures are shown in Fig. S1 a to g. To determine the DP cell population, mRNA expression levels of *Pdgfra, Itga8* and *eGFP* were examined by qRT-PCR (Fig. S1k, l). Cells were sorted with a FACSAria II (BD Biosciences, CA, USA) according to the expression of reporter eGFP and cell surface markers, after gating out dead and Lin^+^ cells. We prepared three or four independent biological replicates and used them for qRT-PCR and RNA-seq.

### qRT-PCR

qRT-PCR was performed as described previously^12^. Primer sequences of the target genes are shown in Supplementary Table 5.

### RNA sequencing, mapping and expression quantification

RNA sequencing and data processing were performed as described previously^12^. Briefly, 10 ng of total RNA samples extracted from FACS-isolated cells were subjected to library preparation using TruSeq Stranded mRNA Sample Prep Kit (Illumina) following the manufacturer’s protocol with minor modification (shortened initial RNA fragmentation to 7 min). We generated three or four biologically independent cDNA libraries for each cell population. The prepared libraries were sequenced using the Rapid Run mode with 80 cycles on the HiSeq1500 (Illumina) followed by trimming low-quality bases and removal of adaptor sequences. The processed reads were mapped to the mm10 mouse genome assembly using the TopHat v2.0.14 with default parameter settings. Gene expression quantification was performed using the Cuffdiff program in the Cufflinks package v2.2.1. Normalized FPKM gene expression values calculated by Cuffduff were used for the following analyses. RNA sequencing and data processing described above were performed at the Laboratory for Phyloinformatics, BDR, RIKEN. Expression data for epidermal populations used in this study were reported in our previous study and deposited in BioProject (PRJNA342736)^12^. RNA-seq data obtained in this study have been submitted to the Sequence Read Archive (SRA) as BioProject: PRJDB9477. RNA-seq read and mapping statistics for the analysed libraries are summarized in Supplementary Table 6.

### Gene expression analysis

To understand the expression patterns of ECM genes, we first compiled a list of ECM genes from the literature^9, 12, 66, 67, 68^, and then defined 281 genes as our matrisome ECM genes (Table S2). To compare gene expression levels among the sorted cell compartments, log2-transformed FPKM values from RNA-seq data were used. For further analysis, genes with low expression (FPKM of less than 3 in all regions) were filtered out and not used. Charts of hierarchical clustering, expression correlation and principal component analysis (PCA) were plotted using Bioconductor R (ver. 3.5.3). For hierarchical clustering, similarity was calculated using the hclust function with Euclidean distance and the complete linkage clustering method. The obtained data were further analysed using the prcomp function for PCA. For gene expression correlation, the cor function with the Spearman method was used. Each ECM gene expression was visualized using the heatmap.2 function.

To elucidate the regional expression of ECM genes, expression levels (average FPKM among replicates) were binarized by setting a threshold (40% value of average FPKM of the epidermal or dermal cell populations that exhibit the highest expression level). Expression values of FPKM less than 3 were always considered to reflect no expression. Then, expressed ECM genes were mapped in a Venn diagram as pie charts, in which the pie size represents the expression levels of the gene. Pie chart graphs were generated using Cytoscape (ver. 3.4.0 with Java 1.8.0).

For Gene Ontology analysis, ECM gene sets were subjected to a statistical overrepresentation test on the PANTHER website (ver. 14.1). Enriched biological process terms (over 40-fold enrichment) were evaluated for their *p*-value and FDR.

### Antibody production

To obtain specific antibody against CRIM1 protein, a Japanese White rabbit was immunized with the recombinant extracellular region of Crim1 protein and raised serum was collected. In detail, a cDNA fragment encoding the extracellular region of mouse *Crim1* (Leu^35^–Asp^939^) was amplified using cDNA derived from E16.5 mouse embryos with the following restriction enzyme site-tagged primer set: forward, GCGGCCCAGCCGGCCCTGGTCTGCCTGCCCTGTG, and reverse, CTCCTCGAGAGAGTCCAGTGATGAGTCTTC. Amplified cDNA was subcloned into the Sfi I-Xho I site of pSecTag2A mammalian expression vector (Invitrogen). The CRIM1 extracellular region was transiently expressed and secreted by 293F cells using ExpiFectamine 293 (Gibco), and purified with a Ni-Sepharose 6 FF column (GE Healthcare, Little Chalfont, UK), following the manufacturer’s protocol. Rabbits were immunized with the purified protein and high-titre serum was obtained (T.K. Craft Corp., Gunma, Japan). Antibody specificity was confirmed by immunostaining using mouse embryonic skin.

Rabbit antiserum to mouse laminin α5 was generated by immunizing rabbits with GST-fused I and II domains of laminin α5 (Lys^2220^–Leu^2459^). The I and II domains were amplified using cDNA derived from E16.5 mouse embryos with the following restriction enzyme site-tagged primer set: forward, CGGGATCCCGTAAACTCCGGAGCCCACCGGGAC, and reverse, GGAATTCCTACTTGTCATCGTCGTCCTTGTAATCCAGGTGCTCTAGGTCCTCC TTAG. Amplified cDNA was subcloned into the EcoR I site of the pGEX-6P-1 expression vector (GE Healthcare). The antigen was expressed in BL21 and purified with a Glutathione Sepharose 4B column (GE Healthcare), following the manufacturer’s protocol. The antibody in the antiserum was affinity-purified with antigen-conjugated CNBr-activated Sepharose 4B. The specificity of the antibody to mouse laminin α5 chain was confirmed by the absence of antibody immunoreactivity to tissue samples from mice with *Lama5* conditional knockout.

### Antibodies

Details of the antibodies used in this study are summarized in Supplementary Table 7.

### Immunohistochemistry and imaging

Whole-mount immunostaining of mouse dorsal skin was performed as described previously^15^. Briefly, mouse skin tissues were dissected and fixed with 4% paraformaldehyde (PFA)/PBS for 1 h at 4°C, and embedded in OCT compound after washing with PBS. For acetone fixation, dissected skin was directly embedded in OCT compound. Skin sections (150 μm thick) were made using a cryostat (Leica, Wetzlar, Germany) and washed with PBS. Acetone fixation was performed by placing skin sections in −30°C acetone for 15 min, followed by acid treatment with 0.1 N HCl/0.1 N KCl for 15 min after washing in PBS. Skin sections were blocked with a blocking buffer (0.5% skim milk/0.25% fish skin gelatin/0.5% Triton X-100/PBS) for 1 h at 4°C, and then incubated with primary antibodies diluted in blocking buffer overnight at 4°C. Skin samples were washed with 0.2% Tween 20/PBS for 4 h and then incubated with secondary antibodies similarly to the primary antibodies. After that, skin samples were stained with DAPI, washed with 0.2% Tween 20/PBS for 4 h at 4°C, and mounted with BABB clearing solution. Images were acquired using Leica TSC SP8. Three-dimensional reconstructed images were produced using Imaris software (Bitplane, Oxford, UK).

### Transmission electron microscopy

Mouse dorsal skin tissues were dissected into 2–3 mm squares and immersed in fixation solution (2% paraformaldehyde/2% glutaraldehyde/0.1 M phosphate buffer). The following steps were performed by Hanaichi Ultrastructure Research Institute (Okazaki, Japan). After washing with 0.1 M phosphate buffer, samples were post-fixed with 2% osmium tetroxide followed by step-dehydration with gradual substitution in higher-concentration ethanol (30%, 50%, 70%, 90% and 100%) and finally 100% propylene oxide. Then, the samples were embedded in epoxy resin Epon812. Ultra-thin sections were cut, stained with uranyl acetate and lead citrate solution, and viewed with a JEM-1200EX (JEOL, Tokyo, Japan) transmission electron microscope at an accelerating voltage of 80 kV.

### Image quantification

All quantification analyses were performed using Fiji software (ver. 2.0.0-rc-69). To calculate ECM protein intensities in the different BM regions, six epidermal regions [interfollicular epidermis (IFE), lower isthmus, upper bulge, mid-bulge, lateral HG and interface region between HG and DP] were specified from HF morphology and their representative immunohistochemical patterns. The target BM regions were manually drawn in the colour split image, and their mean intensities were measured. Relative intensities were calculated as percentile values where the maximum-intensity region was 100. The data were represented as a heatmap chart by Bioconductor R and radar charts by Excel (Microsoft corp., WA, USA). To quantify hemidesmosome-like structures and cellular protrusions, cellular perimeters facing the BM region or space of interest were measured by tracing freehand with a pen on the scale-set images. The lengths were used for calculation of the frequencies of appearance of hemidesmosome-like structures and cellular protrusions. Segmentation of the cellular fraction was manually performed using Fiji software. To quantify dorsal pigmented areas, binarized ROI images were first generated from the individual photos using Fiji software. Image-specific thresholds were determined manually between pigmented and non-pigmented areas from 8-bit greyscale images. Pixels corresponding to the pigmented area were counted using Fiji. A box plot graph was created using Bioconductor R.

### Statistical analysis

Statistical parameters including the numbers of samples and replicates, types of statistical analysis and statistical significance are indicated in the Results, Figures and Figure Legends. \**p*<0.05; \*\**p*<0.01; \*\*\**p*<0.001.

