## Supplementary information for "Mapping the molecular and structural specialization of the skin basement membrane for inter-tissue interactions"

### Supplementary Figure

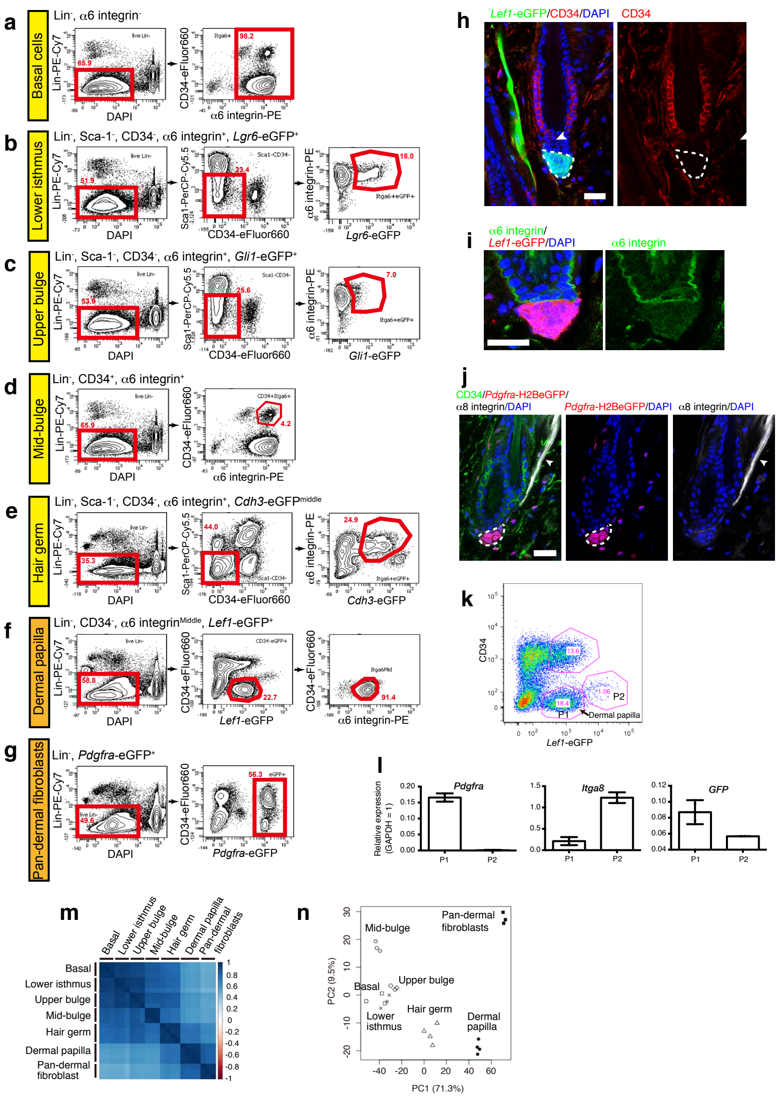


**Supplementary Fig. 1** Isolation of hair follicle epidermal and dermal cell sub-populations. **a–e** FACS-based cell isolation procedures for basal epidermal cells in whole dorsal epidermis (basal cells) (a), lower isthmus (b), upper bulge (c), mid-bulge (d) and HG (e), according to our previous study[11]. Whole dorsal skin from 8-week-old female mice was used. Red polygons indicate sorting gates. **f, g** FACS-based cell isolation procedures for DP (f) and pan-dermal fibroblasts (g). **h** Expression pattern of CD34 in the *Lef1*-eGFP-positive cell populations. Neither DP cells (encircled by dashed line) nor arrector pili muscles (white arrowhead) express CD34. **i** Weak expression of α6 integrin in the *Lef1*-eGFP-positive telogen DP cells. **j** Distinct marker expression in DP and arrector pili muscles. Eight-week-old adult telogen skin of *Pdgfra*-H2BeGFP mice was stained for CD34, GFP, α8 integrin and nuclei (DAPI). DP (encircled by dashed line) cells were CD34$^{-}$/PDGFRA$^{+}$/α8 integrin$^{low}$. Arrector pili muscles were CD34$^{dim}$/PDGFRA$^{-}$/α8 integrin$^{high}$. **k** FACS gate setting for DP cell isolation. Upon Lin$^{+}$ and dead cell depletion, single-cell suspensions of 8-week-old adult telogen dermal tissues of *Lef1*-eGFP mice were plotted based on the signal intensity of CD34 and *Lef1*-eGFP. Two gates (P1 and P2) were used to isolate CD34$^{-}$/*Lef1*-eGFP$^{middle}$ (P1) and CD34$^{middle}$/*Lef1*-eGFP$^{high}$ (P2). **l** Quantitative RT-PCR gene expression analysis of DP and arrector pili muscle marker genes in P1 and P2 cell populations in k. The P1 population showed high *Pdgfra* expression and low *Itga8* expression, consistent with the character of DP. In contrast, the P2 cell population showed no *Pdgfra* expression and high *Itga8* expression, indicating that this cell population is arrector pili muscles. *Lef1*-eGFP expression was detected in both P1 and P2 populations. Data are mean with SD. n = 2 mice. **m** Global gene expression correlation among HF cell populations. Each cell represents overall transcriptomic expression similarity between a pair of samples using Spearman rank correlation coefficient analysis with a range from −1 to 1. **n** Principal component analysis (PCA) of all cell populations analysed in this study. All biological replicates of the same population form close clusters, indicating high reproducibility of our method. Scale bars, 20 μm.

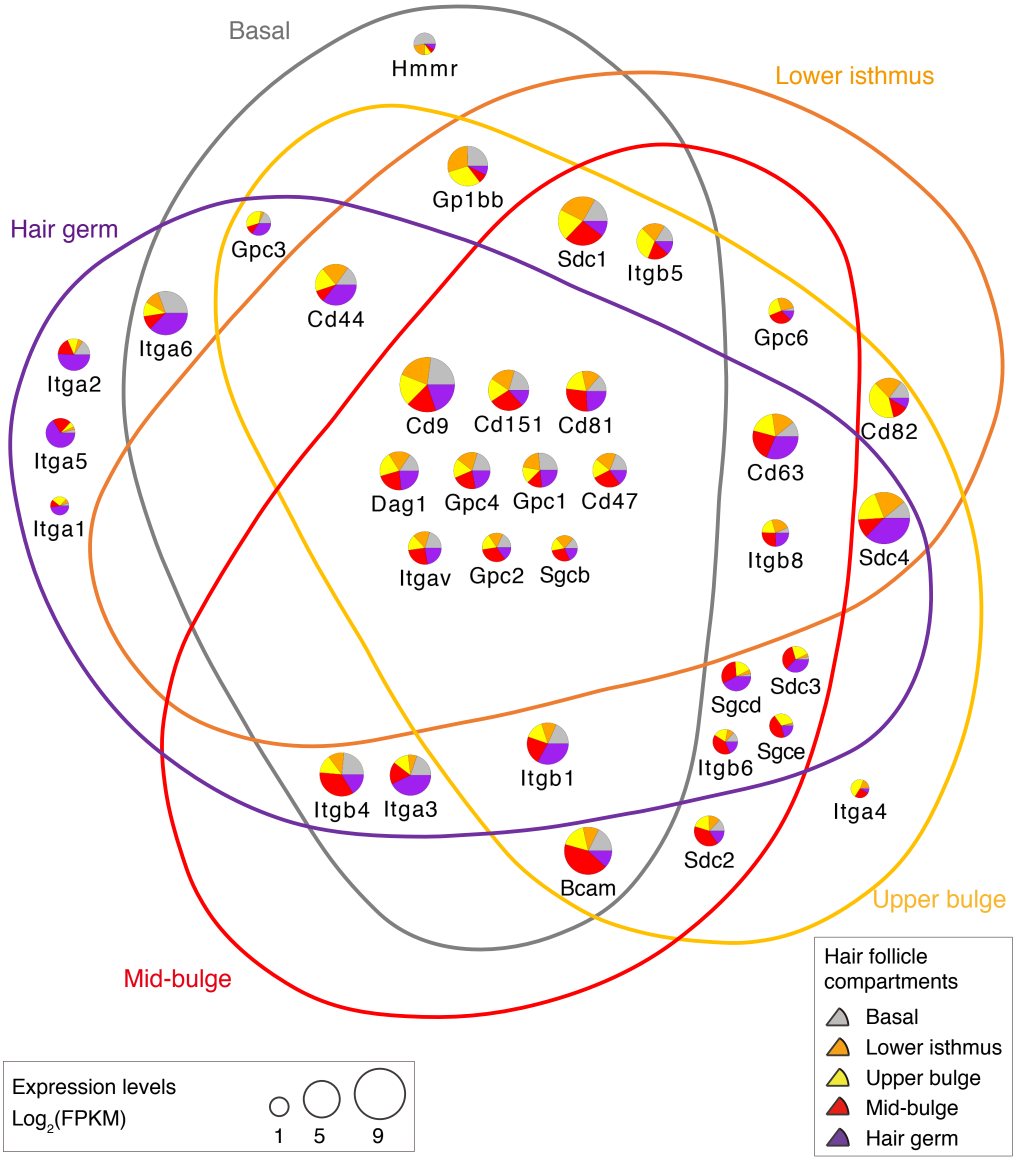


**Supplementary Fig. 2** Venn diagram representing the expression patterns of ECM receptor genes in the epidermal cell populations. This Venn diagram was created in the same way as described in Fig. 3a and Fig. 4a. Tetraspanin genes such as *Cd9*, *Cd151*, *Cd81* and *Cd63* and some glypican genes, *Gpc1* and *Gpc2*, are broadly expressed. While *Itgav* is commonly expressed, other integrin genes are expressed in a region-specific manner. In particular, genes encoding collagen-type integrins, *Itga1* and *Itga2*, and one of the fibronectin receptors, *Itga5*, are highly expressed by the HG cell population.

12

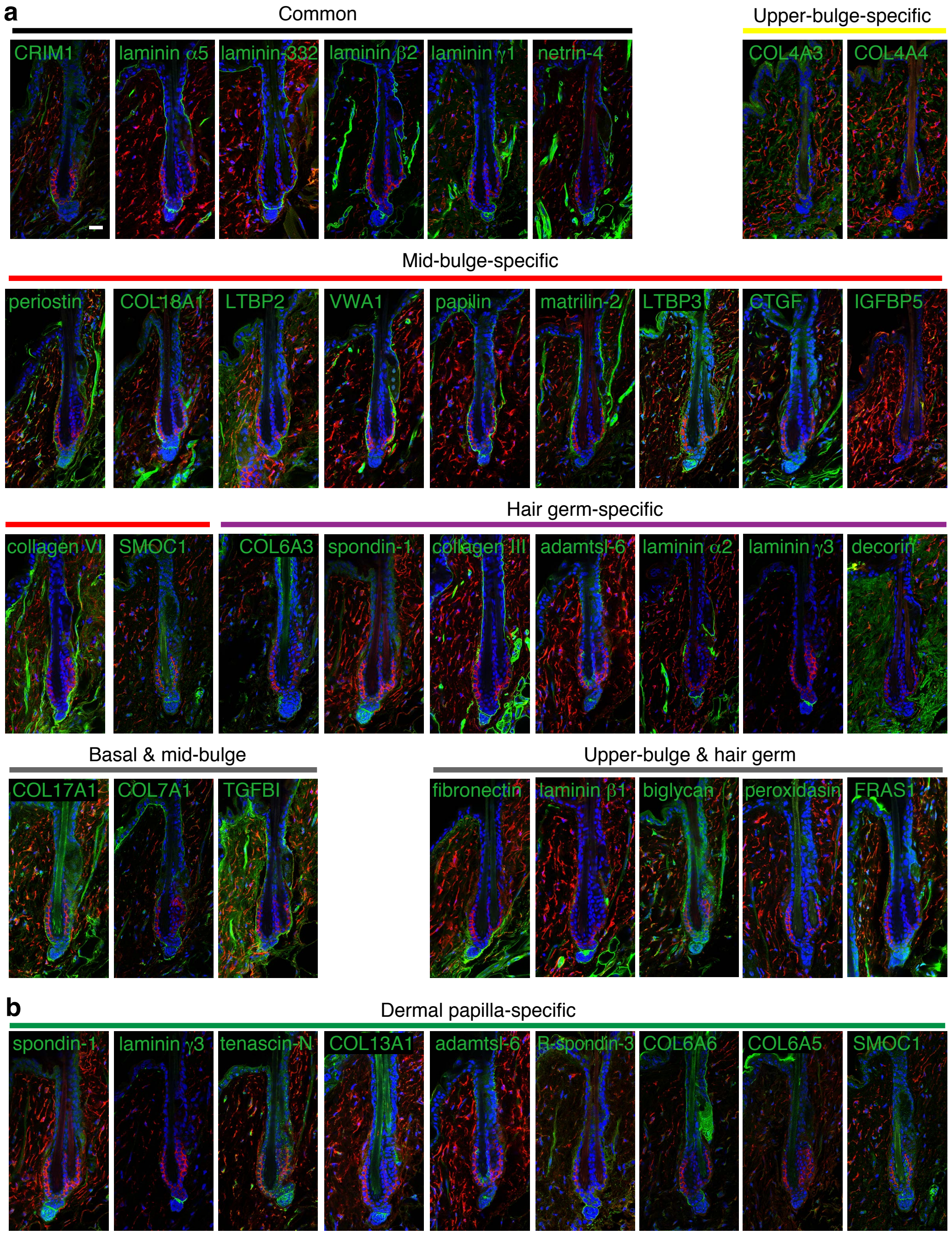


**Supplementary Fig. 3** Immunolocalizations of ECM proteins in the telogen hair follicle. **a, b** Tissue localizations of ECM proteins commonly or region-specifically expressed by the epidermal (a) and dermal populations (b). Individual ECM components (green) were stained with CD34 (red) and DAPI (blue). Their major mRNA sources are indicated at the top of each panel. Scale bar, 20 μm.

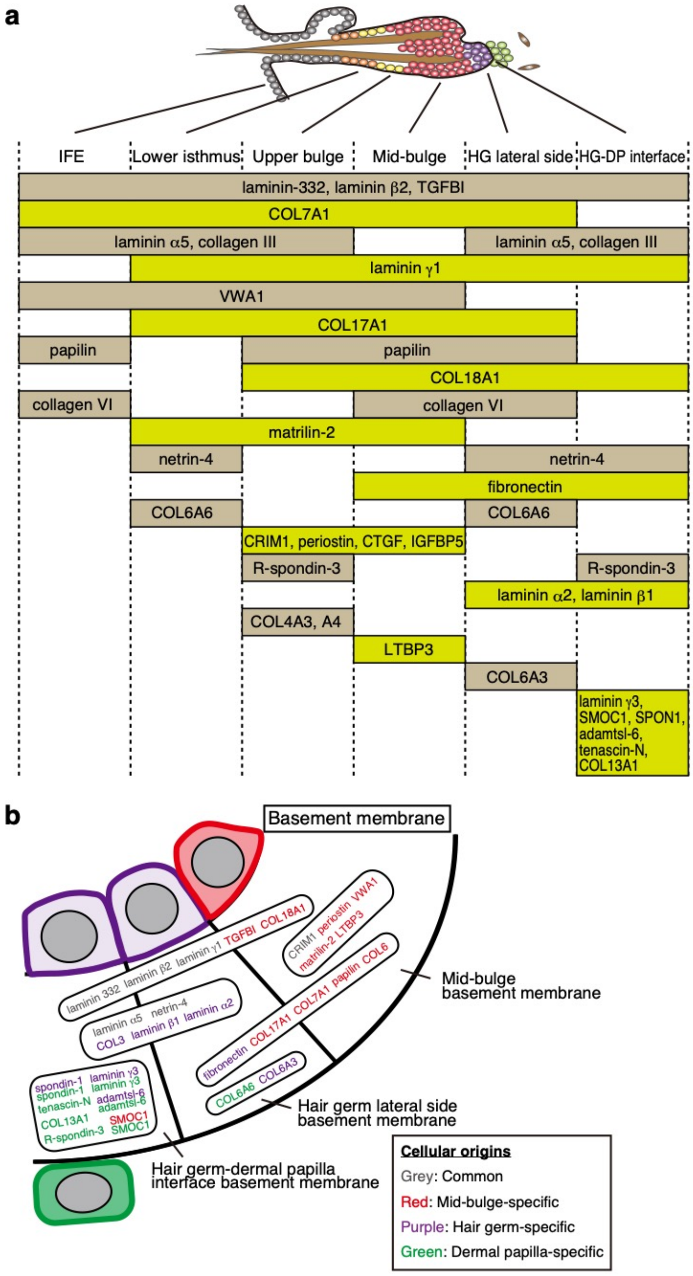


**Supplementary Fig. 4** Graphic display of ECM localizations in telogen hair follicle basement membrane. **a, b** Quantified results of ECM protein deposition shown in Fig. 5b are schematically summarized (a). BM zone just below mid-bulge (red cell) and HG (purple cells) cells is separated into three regions, mid-bulge BM, HG lateral side BM and HG–DP interface BM (b). ECM proteins showing the same deposition patterns (in one specific region or across multiple regions) are grouped in rounded rectangles. Colour-coded proteins indicate their cellular origins (Box).

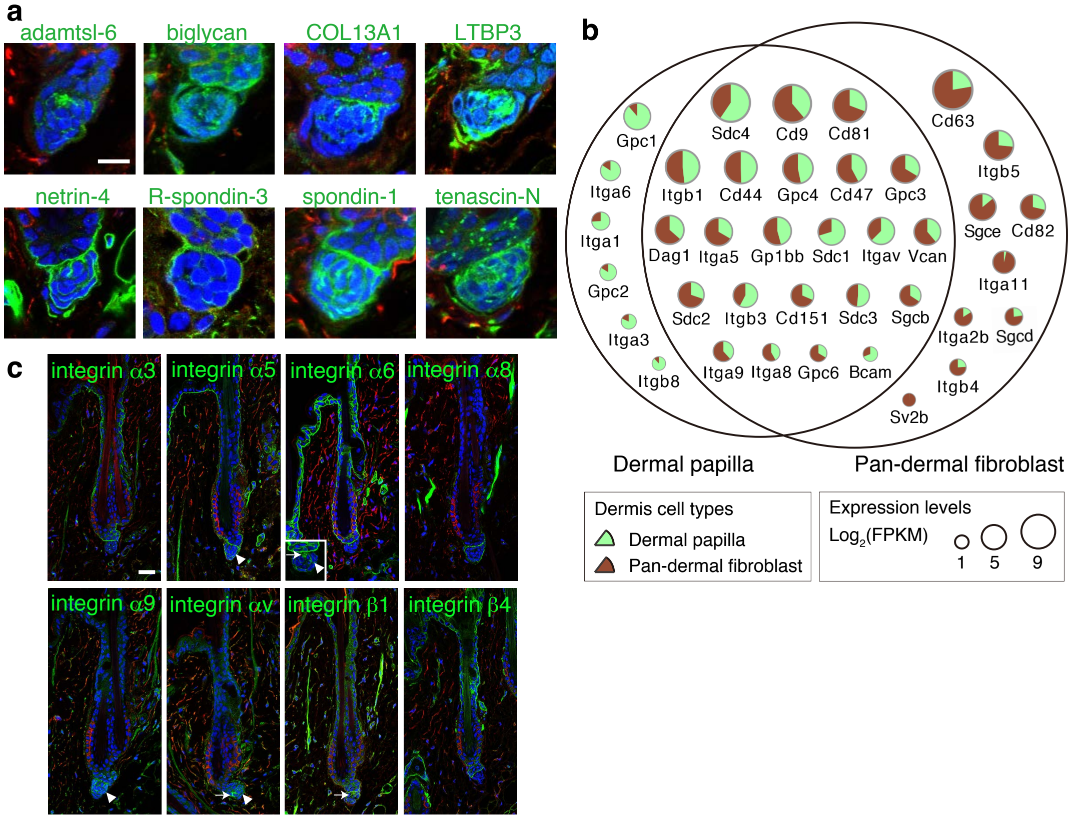


**Supplementary Fig. 5** Tissue localization of ECM and ECM receptor proteins in the hook and mesh basement membranes. **a** Localizations of ECM proteins composing hook and mesh BMs. Images are magnified views of images in Fig. S3. **b** Venn diagram representation of the expression patterns of ECM receptor genes in the dermal cell populations. This Venn diagram was created in the same way as described in Fig. 3a and Fig. 4a. **c** Immunolocalizations of various integrin receptors in the dorsal telogen HFs. Integrins are shown in green with CD34 (red) and nuclear (blue) stains. White arrows and arrowheads indicate hook BM-like localizations and mesh BM-like localizations of integrins, respectively. The inset in the integrin α6 image shows the DP region of a different HF. Scale bars: 10 μm (a) and 20 μm (c).

18

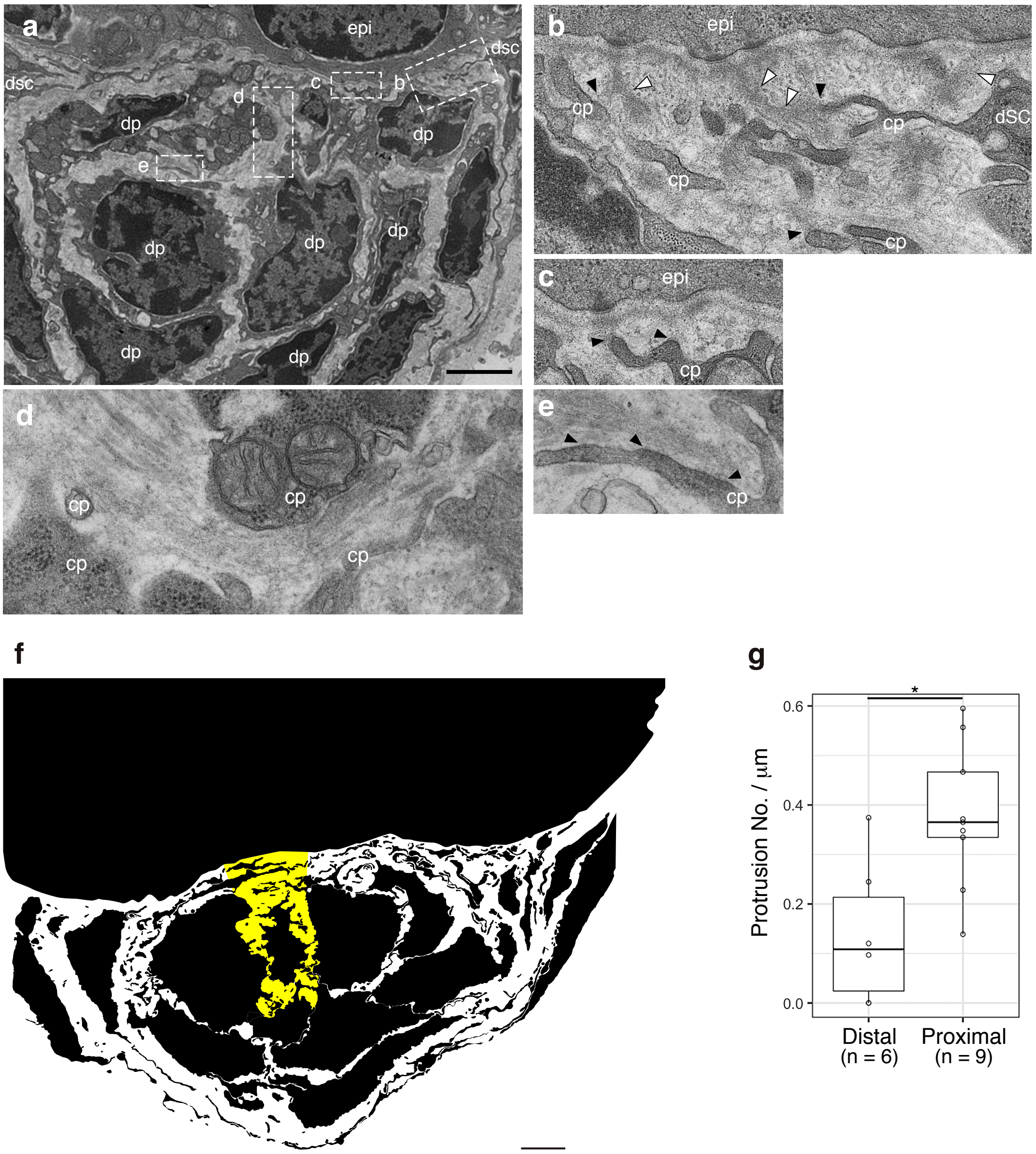


**Supplementary Fig. 6** Cellular protrusions extending toward the basement membrane structures within the dermal papilla. **a** TEM image of the DP. Dotted rectangles indicate the regions of interest magnified in (b–e). **b–e** Magnified views of characteristic ECM structures. Cellular protrusions (cp) interact with protrusions of the interface BM (white arrowheads). Black filled arrowheads indicate the interaction sites. HF dermal stem cells located at the upper edge of DP extended long protrusions toward the interface between HG and DP and appeared to form direct contacts with these BM protrusions and the interface BM (b, c). DP cells extended protrusions toward the hook BM and appeared to form cell adhesion-like structures (d, e). Cellular protrusions have electron-dense membrane structures at cell–BM interfaces (black filled arrowheads in b, c, e). Large extracellular space at the centre of the DP is filled with thick bundles of ECMs (d). **f** Binarized image of cellular fraction and extracellular space in the DP. The cellular fraction is filled with black colour. Large extracellular space at the central region, where the hook BM is located, is marked with yellow. **g** Quantitative comparison of protrusion numbers at the proximal and distal sides to the hook BM. * $p$ <0.05, Mann–Whitney U test. epi, epidermal HG cell; dp, DP cell; dsc, dermal stem cell; cp, cellular protrusion. Scale bars: 2 μm.

19

**Supplementary Table 1.** Gene expression FPKM values of 281 ECM genes in skin cell populations

| Gene Symbol | Basal | Lower isthmus | Upper bulge | Mid-bulge | Hair germ | Dermal papilla | Pan-dermal fibroblasts |
|---|---|---|---|---|---|---|---|
| Lama1 | 5.1 | 0.8 | 0.4 | 0.2 | 0.0 | 0.0 | 0.0 |
| Lama2 | 0.0 | 0.1 | 0.1 | 0.0 | 3.3 | 75.8 | 88.0 |
| Lama3 | 34.3 | 19.8 | 23.3 | 34.1 | 45.9 | 2.3 | 0.7 |
| Lama4 | 1.6 | 4.5 | 3.1 | 1.1 | 2.0 | 24.0 | 81.3 |
| Lama5 | 33.2 | 27.9 | 47.6 | 55.3 | 65.3 | 3.3 | 4.2 |
| Lamb1 | 1.5 | 1.2 | 15.6 | 3.5 | 36.0 | 49.6 | 86.1 |
| Lamb2 | 20.4 | 18.3 | 16.8 | 28.8 | 23.8 | 63.9 | 165.6 |
| Lamb3 | 129.8 | 90.3 | 105.2 | 246.9 | 76.5 | 1.1 | 1.0 |
| Lamc1 | 13.2 | 7.5 | 11.1 | 12.6 | 17.0 | 95.6 | 123.1 |
| Lamc2 | 52.7 | 22.9 | 37.3 | 65.0 | 85.8 | 10.6 | 3.1 |
| Lamc3 | 0.0 | 0.0 | 0.0 | 0.0 | 6.4 | 84.6 | 32.6 |
| Col4a1 | 4.7 | 3.1 | 14.8 | 17.8 | 24.7 | 147.3 | 239.0 |
| Col4a2 | 6.4 | 3.7 | 11.3 | 24.1 | 13.1 | 94.0 | 117.4 |
| Col4a3 | 0.2 | 1.6 | 6.5 | 1.8 | 0.7 | 0.1 | 0.8 |
| Col4a4 | 0.3 | 2.2 | 6.9 | 2.5 | 0.7 | 0.0 | 1.1 |
| Col4a5 | 12.3 | 8.7 | 7.7 | 8.5 | 3.8 | 0.3 | 3.7 |
| Col4a6 | 16.9 | 7.9 | 6.4 | 9.1 | 6.7 | 0.3 | 0.5 |
| Col6a1 | 13.5 | 0.7 | 6.1 | 173.2 | 23.1 | 318.8 | 553.2 |
| Col6a2 | 2.3 | 1.0 | 7.1 | 63.8 | 14.4 | 246.5 | 476.0 |
| Col6a3 | 0.1 | 0.1 | 0.2 | 0.0 | 15.1 | 324.8 | 201.6 |
| Col6a4 | 0.0 | 0.0 | 0.0 | 0.0 | 0.0 | 0.2 | 0.4 |
| Col6a5 | 0.0 | 0.0 | 0.0 | 0.0 | 0.2 | 11.3 | 1.1 |
| Col6a6 | 0.1 | 0.3 | 0.3 | 0.0 | 2.1 | 41.0 | 2.9 |
| Col7a1 | 65.5 | 20.3 | 24.0 | 143.9 | 40.9 | 35.2 | 52.4 |
| Col8a1 | 1.5 | 2.5 | 1.3 | 1.1 | 0.4 | 0.1 | 1.4 |
| Col8a2 | 4.9 | 6.2 | 14.3 | 110.9 | 5.7 | 0.4 | 5.6 |
| Col15a1 | 0.0 | 0.0 | 0.1 | 0.0 | 2.4 | 52.8 | 264.1 |
| Col17a1 | 1086.2 | 517.3 | 518.8 | 2018.0 | 530.7 | 6.4 | 3.0 |
| Col18a1 | 26.0 | 8.5 | 31.4 | 169.2 | 56.9 | 8.5 | 91.5 |
| Col28a1 | 0.0 | 0.0 | 0.0 | 0.0 | 0.2 | 0.4 | 0.5 |
| Nid1 | 0.0 | 0.2 | 0.1 | 0.0 | 1.8 | 33.9 | 73.0 |
| Nid2 | 0.0 | 0.0 | 0.0 | 0.0 | 0.9 | 7.7 | 4.0 |
| Hspg2 | 4.7 | 5.2 | 15.3 | 20.7 | 23.1 | 28.3 | 112.3 |
| Acan | 0.0 | 0.0 | 0.0 | 0.0 | 0.0 | 2.4 | 0.6 |

| | | | | | | | |
|---|---|---|---|---|---|---|---|
| Agrn | 89.8 | 60.0 | 111.0 | 248.0 | 128.8 | 20.4 | 9.1 |
| Amelx | 0.0 | 0.0 | 0.0 | 0.0 | 0.0 | 0.0 | 0.0 |
| Colq | 0.1 | 0.0 | 0.0 | 0.3 | 0.0 | 0.0 | 5.4 |
| Efemp1 | 27.6 | 8.4 | 16.8 | 3.9 | 18.0 | 6.6 | 857.6 |
| Egfl6 | 7.6 | 8.7 | 25.0 | 40.7 | 6.1 | 0.4 | 0.1 |
| Egflam | 0.1 | 0.1 | 0.6 | 2.8 | 1.2 | 23.9 | 21.0 |
| Emilin3 | 0.3 | 0.3 | 0.7 | 7.0 | 0.3 | 0.1 | 0.3 |
| Fbln1 | 3.6 | 12.4 | 47.2 | 50.4 | 42.7 | 25.0 | 231.1 |
| Fras1 | 0.1 | 0.1 | 3.7 | 0.1 | 7.2 | 0.5 | 0.2 |
| Frem1 | 0.0 | 0.0 | 0.9 | 0.0 | 1.1 | 7.4 | 9.8 |
| Frem2 | 11.0 | 4.1 | 8.1 | 7.2 | 5.6 | 0.3 | 0.1 |
| Frem3 | 0.0 | 0.2 | 0.0 | 0.0 | 0.0 | 0.0 | 0.0 |
| Hmcn1 | 2.4 | 2.4 | 7.2 | 25.4 | 20.6 | 1.7 | 7.7 |
| Matn2 | 6.5 | 6.5 | 5.9 | 20.6 | 5.8 | 15.1 | 33.3 |
| Mmrn2 | 0.0 | 0.0 | 0.0 | 0.1 | 0.0 | 0.1 | 0.1 |
| Npnt | 10.9 | 3.7 | 27.9 | 67.9 | 24.2 | 2.1 | 0.2 |
| Ntn1 | 0.0 | 0.0 | 0.0 | 0.1 | 0.2 | 14.1 | 103.3 |
| Ntn3 | 0.3 | 0.6 | 1.9 | 1.8 | 1.4 | 4.2 | 3.6 |
| Ntn4 | 3.1 | 3.3 | 3.4 | 4.6 | 4.4 | 1.2 | 19.1 |
| Ntn5 | 0.0 | 0.0 | 0.0 | 0.0 | 0.2 | 0.0 | 10.4 |
| Ntng1 | 0.0 | 0.0 | 0.2 | 0.0 | 0.1 | 2.1 | 4.1 |
| Papln | 5.8 | 1.5 | 3.1 | 22.8 | 3.9 | 0.3 | 0.4 |
| Smoc1 | 0.9 | 0.2 | 0.8 | 5.8 | 2.7 | 5.4 | 0.9 |
| Smoc2 | 145.2 | 83.3 | 60.1 | 14.1 | 41.9 | 62.4 | 78.7 |
| Sparc | 31.0 | 3.2 | 27.3 | 260.1 | 35.6 | 212.2 | 8352.8 |
| Tgfbi | 378.0 | 185.4 | 129.0 | 538.7 | 79.4 | 2.9 | 1253.0 |
| Thbs2 | 34.7 | 20.5 | 10.0 | 14.1 | 6.8 | 288.6 | 574.7 |
| Thbs4 | 0.0 | 0.0 | 0.0 | 0.0 | 0.0 | 0.0 | 25.4 |
| Tinag | 0.0 | 0.0 | 0.0 | 0.0 | 0.0 | 0.0 | 0.1 |
| Tinagl1 | 1.7 | 2.6 | 1.8 | 1.3 | 5.7 | 2.8 | 1.5 |
| Tnc | 1.8 | 0.9 | 5.5 | 19.5 | 38.5 | 96.9 | 39.2 |
| Vwa1 | 6.2 | 6.3 | 5.9 | 40.1 | 2.0 | 0.5 | 1.1 |
| Vwa2 | 12.7 | 38.1 | 111.9 | 122.1 | 20.3 | 0.4 | 0.1 |
| Col1a1 | 1.0 | 1.4 | 3.5 | 2.8 | 5.3 | 75.4 | 13411.7 |
| Col1a2 | 1.4 | 1.3 | 9.3 | 7.6 | 8.1 | 92.9 | 12614.0 |
| Col2a1 | 0.0 | 0.0 | 0.1 | 0.1 | 0.5 | 0.1 | 0.1 |
| Col3a1 | 0.3 | 0.5 | 0.6 | 0.0 | 5.9 | 155.5 | 5390.8 |

| | | | | | | | |
|---|---|---|---|---|---|---|---|
| Col5a1 | 0.3 | 0.8 | 4.3 | 3.8 | 2.8 | 30.7 | 261.8 |
| Col5a2 | 1.1 | 5.0 | 14.4 | 19.1 | 8.8 | 67.7 | 414.4 |
| Col5a3 | 0.1 | 0.0 | 0.1 | 0.2 | 0.3 | 5.1 | 99.4 |
| Col9a1 | 0.0 | 0.0 | 0.0 | 0.0 | 0.0 | 0.0 | 0.0 |
| Col9a2 | 0.0 | 0.3 | 1.6 | 0.2 | 0.1 | 0.2 | 0.1 |
| Col9a3 | 0.1 | 0.5 | 0.4 | 0.5 | 0.2 | 0.2 | 0.5 |
| Col10a1 | 0.0 | 0.0 | 0.0 | 0.0 | 0.0 | 0.1 | 0.1 |
| Col11a1 | 0.0 | 0.0 | 0.0 | 0.0 | 0.0 | 2.3 | 16.2 |
| Col11a2 | 0.4 | 0.4 | 0.3 | 0.2 | 0.3 | 0.6 | 0.4 |
| Col12a1 | 4.0 | 1.0 | 1.8 | 21.4 | 4.4 | 11.7 | 11.3 |
| Col13a1 | 0.0 | 0.0 | 0.0 | 0.0 | 1.8 | 38.2 | 8.2 |
| Col14a1 | 0.1 | 0.1 | 2.8 | 0.4 | 2.3 | 5.8 | 25.2 |
| Col16a1 | 35.9 | 6.1 | 7.8 | 20.2 | 15.0 | 117.9 | 393.0 |
| Col19a1 | 0.0 | 0.0 | 0.0 | 0.0 | 0.0 | 0.0 | 0.0 |
| Col20a1 | 2.5 | 3.4 | 3.8 | 3.9 | 4.2 | 1.1 | 2.2 |
| Col22a1 | 0.0 | 0.0 | 0.0 | 0.0 | 0.0 | 0.0 | 0.1 |
| Col23a1 | 25.5 | 4.7 | 0.3 | 0.6 | 5.9 | 111.8 | 23.3 |
| Col24a1 | 0.0 | 0.0 | 0.1 | 0.1 | 0.0 | 0.3 | 2.7 |
| Col25a1 | 0.0 | 0.0 | 0.5 | 0.0 | 0.4 | 0.1 | 0.1 |
| Col26a1 | 0.0 | 0.0 | 0.0 | 0.0 | 0.0 | 0.2 | 0.3 |
| Col27a1 | 0.0 | 0.1 | 1.2 | 0.1 | 1.2 | 10.7 | 4.0 |
| Aspn | 4.5 | 50.3 | 117.9 | 47.5 | 21.3 | 129.5 | 336.3 |
| Bcan | 0.0 | 0.0 | 0.0 | 0.0 | 0.0 | 0.1 | 0.0 |
| Bgn | 1.2 | 0.9 | 19.5 | 5.3 | 26.8 | 276.1 | 2292.9 |
| Chad | 0.7 | 1.0 | 0.9 | 1.4 | 0.9 | 0.3 | 0.7 |
| Chadl | 0.8 | 0.7 | 1.3 | 3.6 | 0.8 | 0.6 | 2.0 |
| Dcn | 1.5 | 1.7 | 2.5 | 0.0 | 6.7 | 141.1 | 20609.5 |
| Epyc | 0.0 | 0.0 | 0.0 | 0.0 | 0.0 | 0.0 | 0.2 |
| Esm1 | 0.0 | 0.0 | 0.0 | 0.0 | 0.1 | 2.0 | 0.9 |
| Fmod | 0.2 | 2.2 | 4.4 | 1.8 | 1.1 | 1.4 | 606.6 |
| Hapln1 | 0.0 | 0.0 | 0.0 | 0.0 | 0.0 | 0.9 | 0.8 |
| Hapln2 | 0.0 | 0.5 | 0.5 | 0.1 | 0.3 | 0.2 | 0.0 |
| Hapln3 | 0.0 | 0.0 | 0.1 | 0.0 | 0.1 | 1.3 | 2.2 |
| Hapln4 | 0.0 | 0.0 | 0.0 | 0.0 | 0.0 | 0.0 | 2.1 |
| Impg1 | 0.0 | 0.0 | 0.0 | 0.0 | 0.0 | 0.0 | 0.0 |
| Impg2 | 0.1 | 0.0 | 0.0 | 0.0 | 0.1 | 0.1 | 0.6 |
| Kera | 0.0 | 0.0 | 0.0 | 0.0 | 0.2 | 0.3 | 61.1 |

| | | | | | | | |
|---|---|---|---|---|---|---|---|
| Lum | 0.1 | 0.2 | 0.2 | 0.0 | 1.1 | 18.9 | 3060.6 |
| Ncan | 0.0 | 0.0 | 0.0 | 0.0 | 0.0 | 0.0 | 0.0 |
| Nepn | 0.0 | 0.0 | 0.0 | 0.0 | 0.0 | 0.0 | 0.0 |
| Nyx | 0.1 | 0.1 | 0.0 | 0.1 | 0.0 | 0.0 | 0.2 |
| Ogn | 0.0 | 0.0 | 0.0 | 0.0 | 0.0 | 1.8 | 278.6 |
| Omd | 2.3 | 2.4 | 2.3 | 2.0 | 10.9 | 19.3 | 24.9 |
| Optc | 0.5 | 0.7 | 0.7 | 0.3 | 0.4 | 0.4 | 0.6 |
| Podn | 0.4 | 1.6 | 2.6 | 1.3 | 3.9 | 11.8 | 85.2 |
| Podnl1 | 0.0 | 0.0 | 0.0 | 0.0 | 0.0 | 0.5 | 0.1 |
| Prelp | 0.2 | 0.4 | 0.9 | 0.6 | 0.7 | 7.8 | 292.6 |
| Prg2 | 0.0 | 0.0 | 0.0 | 0.0 | 0.0 | 0.1 | 0.0 |
| Prg3 | 0.0 | 0.0 | 0.0 | 0.0 | 0.0 | 0.0 | 0.0 |
| Prg4 | 1.1 | 1.0 | 0.9 | 0.8 | 0.4 | 1.9 | 2.6 |
| Spock1 | 0.0 | 0.1 | 0.1 | 1.0 | 1.5 | 41.7 | 3.5 |
| Spock2 | 2.4 | 3.7 | 2.4 | 4.8 | 6.8 | 1.4 | 14.4 |
| Spock3 | 0.0 | 0.0 | 0.0 | 0.0 | 0.1 | 2.0 | 0.0 |
| Srgn | 0.0 | 0.7 | 0.3 | 0.2 | 4.8 | 76.9 | 12.5 |
| Vcan | 0.0 | 0.0 | 0.2 | 0.0 | 1.5 | 32.7 | 51.4 |
| 5430419D17Rik | 0.0 | 0.0 | 0.0 | 0.0 | 0.0 | 0.0 | 0.0 |
| Abi3bp | 12.5 | 10.8 | 78.3 | 7.2 | 116.9 | 13.0 | 140.1 |
| Adamtsl1 | 0.5 | 0.2 | 1.4 | 0.2 | 1.3 | 0.1 | 16.8 |
| Adamtsl2 | 0.0 | 0.0 | 0.1 | 0.1 | 1.5 | 16.4 | 8.5 |
| Adamtsl3 | 0.0 | 0.0 | 0.0 | 0.5 | 0.0 | 0.0 | 0.8 |
| Adamtsl4 | 8.5 | 8.0 | 16.2 | 118.3 | 6.8 | 2.3 | 23.0 |
| Adamtsl5 | 20.9 | 7.6 | 7.3 | 22.2 | 14.5 | 34.7 | 127.7 |
| Thsd4 | 2.9 | 0.4 | 1.2 | 0.2 | 5.8 | 38.6 | 4.0 |
| Adipoq | 0.0 | 0.0 | 0.0 | 0.0 | 0.2 | 0.1 | 0.3 |
| Aebp1 | 0.8 | 1.6 | 4.2 | 5.8 | 4.2 | 6.2 | 1058.1 |
| Ambn | 0.0 | 0.0 | 0.0 | 0.0 | 0.0 | 0.0 | 0.0 |
| AW551984 | 0.0 | 0.0 | 0.0 | 0.0 | 0.1 | 0.1 | 8.1 |
| Bglap2 | 0.0 | 0.1 | 0.0 | 0.0 | 0.0 | 0.0 | 0.0 |
| Bglap3 | 0.5 | 1.7 | 1.0 | 0.9 | 0.9 | 0.0 | 0.2 |
| Bmper | 0.0 | 0.1 | 0.3 | 0.6 | 0.7 | 0.2 | 0.2 |
| Bsph1 | 0.0 | 0.0 | 0.0 | 0.0 | 0.0 | 0.0 | 0.0 |
| Bsph2 | 0.0 | 0.0 | 0.0 | 0.0 | 0.0 | 0.0 | 0.0 |
| Cdcp2 | 0.0 | 0.0 | 0.0 | 0.0 | 0.0 | 0.0 | 0.0 |
| Cilp | 0.2 | 0.1 | 1.1 | 2.1 | 8.1 | 165.6 | 291.1 |

| | | | | | | | |
|---|---|---|---|---|---|---|---|
| Cilp2 | 0.2 | 0.6 | 0.7 | 0.7 | 1.3 | 0.2 | 0.4 |
| Coch | 0.1 | 0.3 | 0.4 | 0.1 | 2.5 | 36.7 | 2267.4 |
| Comp | 0.1 | 0.1 | 0.0 | 0.1 | 0.1 | 0.1 | 0.1 |
| Creld1 | 4.6 | 5.3 | 3.0 | 3.4 | 3.0 | 3.9 | 18.8 |
| Creld2 | 13.7 | 23.9 | 18.3 | 18.6 | 12.1 | 18.7 | 34.2 |
| Crim1 | 30.7 | 31.3 | 37.3 | 72.4 | 30.0 | 15.0 | 11.5 |
| Crispld1 | 0.4 | 2.5 | 10.3 | 8.5 | 2.6 | 17.0 | 2.9 |
| Crispld2 | 3.0 | 0.7 | 0.1 | 0.0 | 12.7 | 191.8 | 181.5 |
| Ctgf | 40.6 | 140.9 | 244.4 | 713.5 | 103.0 | 9.7 | 192.6 |
| Cthrc1 | 0.0 | 0.1 | 0.1 | 0.5 | 0.1 | 0.3 | 3.2 |
| Cyr61 | 575.8 | 185.8 | 335.7 | 898.8 | 1143.7 | 1990.1 | 1505.4 |
| Ddx26b | 17.2 | 12.0 | 21.5 | 21.4 | 43.4 | 52.4 | 16.3 |
| Dmbt1 | 0.0 | 0.0 | 0.0 | 0.0 | 0.0 | 0.0 | 0.0 |
| Dmp1 | 0.0 | 0.0 | 0.0 | 0.0 | 0.0 | 0.0 | 0.0 |
| Dpt | 0.1 | 0.1 | 0.1 | 0.0 | 8.6 | 161.4 | 986.6 |
| Dspp | 0.0 | 0.0 | 0.0 | 0.0 | 0.0 | 0.0 | 0.0 |
| Ecm1 | 80.3 | 56.8 | 92.9 | 613.1 | 79.4 | 84.7 | 350.3 |
| Ecm2 | 0.0 | 0.1 | 0.1 | 0.1 | 0.6 | 10.9 | 353.8 |
| Edil3 | 0.0 | 0.0 | 0.0 | 0.0 | 0.3 | 5.4 | 0.4 |
| Efemp2 | 1.0 | 0.8 | 6.7 | 11.4 | 10.8 | 37.1 | 82.9 |
| Egfem1 | 0.0 | 0.0 | 0.0 | 0.0 | 0.0 | 0.0 | 0.0 |
| Eln | 0.1 | 0.0 | 0.4 | 0.1 | 0.8 | 3.9 | 463.8 |
| Emid1 | 0.4 | 1.9 | 4.6 | 4.7 | 0.4 | 0.8 | 40.2 |
| Emilin1 | 0.0 | 0.0 | 0.0 | 0.0 | 0.9 | 11.3 | 18.6 |
| Emilin2 | 0.0 | 0.0 | 0.0 | 0.0 | 1.1 | 12.7 | 60.9 |
| Fbln2 | 0.6 | 0.3 | 2.6 | 9.2 | 4.1 | 41.5 | 302.3 |
| Fbln5 | 0.0 | 0.0 | 0.1 | 0.1 | 0.3 | 6.5 | 95.2 |
| Fbln7 | 0.6 | 2.8 | 0.8 | 0.4 | 0.4 | 3.1 | 12.8 |
| Fbn1 | 0.0 | 0.0 | 0.1 | 0.0 | 1.1 | 24.9 | 311.8 |
| Fbn2 | 0.1 | 0.7 | 0.8 | 0.2 | 0.1 | 0.8 | 1.5 |
| Fga | 0.0 | 0.0 | 0.0 | 0.0 | 0.0 | 0.0 | 0.0 |
| Fgb | 0.0 | 0.0 | 0.0 | 0.0 | 0.0 | 0.0 | 0.0 |
| Fgg | 0.0 | 0.0 | 0.4 | 0.0 | 3.1 | 0.0 | 0.0 |
| Fgl1 | 0.2 | 0.2 | 1.0 | 1.1 | 1.0 | 0.2 | 0.3 |
| Fgl2 | 0.1 | 0.1 | 0.3 | 0.1 | 6.9 | 61.2 | 1034.3 |
| Fn1 | 2.5 | 1.2 | 24.3 | 1.7 | 56.1 | 6.5 | 679.7 |
| Fndc1 | 0.2 | 0.1 | 0.2 | 0.3 | 0.2 | 1.0 | 105.4 |

| | | | | | | | |
|---|---|---|---|---|---|---|---|
| Fndc7 | 0.0 | 0.0 | 0.0 | 0.0 | 0.0 | 0.0 | 0.1 |
| Fndc8 | 0.1 | 0.1 | 0.1 | 0.1 | 0.0 | 0.2 | 0.0 |
| Gas6 | 179.1 | 139.5 | 125.0 | 174.8 | 57.7 | 292.2 | 604.6 |
| Gldn | 0.0 | 0.0 | 0.0 | 0.0 | 1.5 | 47.6 | 74.3 |
| Ibsp | 0.0 | 0.0 | 0.0 | 0.0 | 0.0 | 0.0 | 0.0 |
| Igfals | 1.0 | 0.5 | 2.6 | 2.9 | 3.2 | 0.5 | 3.1 |
| Igfbp1 | 0.0 | 0.0 | 0.0 | 0.1 | 0.0 | 0.0 | 0.2 |
| Igfbp2 | 2.3 | 17.3 | 1.2 | 0.6 | 2.6 | 3.1 | 547.3 |
| Igfbp3 | 343.5 | 186.8 | 90.6 | 179.1 | 125.9 | 1667.3 | 370.5 |
| Igfbp4 | 16.7 | 112.5 | 104.9 | 89.9 | 24.0 | 13.9 | 62.6 |
| Igfbp5 | 1.7 | 0.2 | 5.6 | 45.9 | 1.1 | 5.3 | 989.0 |
| Igfbp6 | 1.9 | 1.7 | 2.3 | 42.6 | 3.7 | 34.9 | 603.7 |
| Igfbp7 | 30.3 | 20.2 | 74.0 | 156.0 | 20.4 | 373.9 | 2016.8 |
| Igfbpl1 | 0.0 | 0.0 | 0.0 | 0.0 | 0.0 | 0.0 | 0.0 |
| Igsf10 | 0.4 | 0.3 | 1.1 | 2.7 | 4.1 | 25.0 | 70.4 |
| Kcp | 0.0 | 0.0 | 0.0 | 0.0 | 0.2 | 2.9 | 7.2 |
| Lgi1 | 0.0 | 0.0 | 0.0 | 0.0 | 0.0 | 0.0 | 4.4 |
| Lgi2 | 0.0 | 0.1 | 0.0 | 0.0 | 0.0 | 0.7 | 0.2 |
| Lgi3 | 0.2 | 0.1 | 0.1 | 0.1 | 0.0 | 0.1 | 1.4 |
| Lgi4 | 0.8 | 1.6 | 1.3 | 2.2 | 1.0 | 1.4 | 22.3 |
| Lrg1 | 0.1 | 0.0 | 0.0 | 0.0 | 0.3 | 0.2 | 0.2 |
| Ltbp1 | 2.9 | 5.3 | 17.0 | 18.9 | 8.6 | 69.5 | 76.3 |
| Ltbp2 | 4.9 | 1.1 | 5.0 | 112.7 | 14.1 | 3.7 | 26.3 |
| Ltbp3 | 24.1 | 32.5 | 44.3 | 112.7 | 41.1 | 31.0 | 62.6 |
| Ltbp4 | 76.9 | 34.4 | 70.6 | 162.7 | 98.3 | 47.1 | 486.2 |
| Matn1 | 0.0 | 0.0 | 0.0 | 0.0 | 0.0 | 3.1 | 0.0 |
| Matn3 | 0.0 | 0.0 | 0.0 | 0.0 | 0.0 | 0.0 | 0.0 |
| Matn4 | 0.0 | 0.3 | 0.4 | 0.1 | 0.6 | 0.8 | 5.8 |
| Mepe | 0.0 | 0.0 | 0.0 | 0.0 | 0.0 | 0.0 | 0.0 |
| Mfap1a | 4.6 | 3.9 | 3.7 | 7.6 | 3.4 | 7.5 | 4.9 |
| Mfap1b | 0.6 | 1.0 | 0.9 | 1.3 | 0.7 | 1.6 | 1.2 |
| Mfap2 | 0.7 | 1.5 | 33.4 | 3.7 | 27.7 | 55.5 | 289.5 |
| Mfap3 | 7.1 | 5.0 | 4.3 | 6.8 | 3.0 | 5.9 | 4.7 |
| Mfap4 | 0.2 | 0.2 | 0.1 | 0.0 | 1.2 | 49.6 | 2040.6 |
| Mfap5 | 0.0 | 0.0 | 0.0 | 0.0 | 0.0 | 1.6 | 157.8 |
| Mfge8 | 2.8 | 3.8 | 5.9 | 59.7 | 21.1 | 56.6 | 155.7 |
| Mgp | 3.2 | 11.2 | 24.7 | 13.2 | 54.2 | 6.1 | 130.8 |

| | | | | | | | |
|---|---|---|---|---|---|---|---|
| Mmrn1 | 0.0 | 0.0 | 0.0 | 0.0 | 0.0 | 0.1 | 0.0 |
| Ndnf | 0.1 | 0.0 | 0.0 | 0.0 | 1.5 | 18.8 | 3.7 |
| Nell1 | 0.0 | 0.0 | 0.0 | 0.0 | 0.1 | 0.4 | 0.4 |
| Nell2 | 0.0 | 0.0 | 0.3 | 0.0 | 0.2 | 0.0 | 0.1 |
| Nov | 0.0 | 0.0 | 0.1 | 0.0 | 4.4 | 86.1 | 143.5 |
| Ntng2 | 4.1 | 1.8 | 0.2 | 0.4 | 1.0 | 13.9 | 16.0 |
| Oit3 | 0.0 | 0.0 | 0.0 | 0.0 | 0.0 | 0.1 | 0.2 |
| Otog | 0.0 | 0.0 | 0.0 | 0.0 | 0.0 | 0.0 | 0.0 |
| Otogl | 0.1 | 0.4 | 0.1 | 0.0 | 0.0 | 0.2 | 0.1 |
| Otol1 | 0.0 | 0.0 | 0.0 | 0.0 | 0.0 | 0.0 | 0.0 |
| Pcolce | 0.8 | 0.5 | 0.2 | 2.4 | 4.9 | 104.5 | 826.7 |
| Pcolce2 | 0.5 | 0.1 | 1.6 | 11.9 | 1.4 | 1.4 | 503.9 |
| Postn | 133.6 | 230.7 | 553.6 | 1696.8 | 142.9 | 67.5 | 310.9 |
| Pxdn | 0.8 | 1.4 | 8.1 | 2.9 | 10.0 | 15.9 | 29.1 |
| Reln | 0.0 | 0.0 | 0.0 | 0.0 | 0.1 | 0.1 | 0.2 |
| Rspo1 | 0.0 | 0.0 | 0.4 | 0.7 | 5.3 | 53.6 | 83.1 |
| Rspo2 | 0.0 | 0.0 | 0.0 | 0.0 | 0.1 | 3.3 | 0.4 |
| Rspo3 | 8.0 | 0.8 | 0.9 | 0.2 | 0.9 | 15.2 | 2.6 |
| Rspo4 | 0.0 | 0.0 | 0.0 | 0.0 | 0.3 | 5.6 | 25.3 |
| Sbspon | 0.0 | 0.1 | 0.7 | 0.1 | 0.1 | 0.9 | 0.2 |
| Slamf6 | 0.0 | 0.0 | 0.0 | 0.0 | 0.0 | 0.0 | 0.0 |
| Slit1 | 0.2 | 0.2 | 0.1 | 0.1 | 0.1 | 0.0 | 0.1 |
| Slit2 | 0.0 | 0.0 | 0.0 | 0.0 | 0.9 | 27.1 | 56.0 |
| Slit3 | 2.8 | 5.1 | 1.7 | 6.5 | 6.0 | 66.6 | 205.3 |
| Sned1 | 0.3 | 0.1 | 0.4 | 1.9 | 6.3 | 43.4 | 56.2 |
| Sparcl1 | 0.1 | 0.1 | 0.1 | 0.0 | 1.2 | 22.3 | 213.5 |
| Spon1 | 0.0 | 0.0 | 0.1 | 0.0 | 11.2 | 223.2 | 28.4 |
| Spon2 | 2.3 | 4.4 | 8.9 | 23.2 | 1.5 | 15.7 | 136.6 |
| Spp1 | 0.1 | 0.1 | 0.0 | 0.0 | 15.7 | 645.2 | 31.9 |
| Srpx | 0.0 | 0.0 | 0.5 | 0.5 | 0.7 | 8.0 | 222.5 |
| Srpx2 | 0.0 | 0.0 | 0.0 | 0.0 | 0.0 | 1.2 | 29.2 |
| Sspo | 0.0 | 0.0 | 0.1 | 0.2 | 0.2 | 0.0 | 0.1 |
| Svep1 | 0.0 | 0.0 | 0.0 | 0.0 | 0.2 | 3.9 | 52.2 |
| Tecta | 0.0 | 0.0 | 0.0 | 0.0 | 0.0 | 0.1 | 0.0 |
| Tectb | 0.0 | 0.0 | 0.0 | 0.0 | 0.0 | 0.0 | 0.0 |
| Thbs1 | 390.4 | 115.0 | 225.4 | 142.2 | 1036.7 | 813.1 | 622.7 |
| Thbs3 | 5.6 | 1.6 | 5.5 | 11.9 | 16.0 | 31.7 | 89.9 |

| | | | | | | | |
|---|---|---|---|---|---|---|---|
| Tnfaip6 | 0.3 | 0.3 | 0.5 | 0.8 | 57.9 | 523.4 | 2814.0 |
| Tnn | 0.1 | 0.1 | 0.1 | 0.1 | 0.5 | 57.6 | 20.1 |
| Tnr | 0.0 | 0.0 | 0.0 | 0.0 | 0.0 | 0.0 | 0.0 |
| Tnxb | 34.7 | 9.6 | 1.2 | 3.8 | 3.4 | 1.0 | 295.3 |
| Tsku | 0.7 | 1.5 | 4.8 | 5.3 | 3.4 | 2.8 | 7.4 |
| Vit | 0.5 | 0.3 | 2.0 | 7.2 | 0.2 | 0.9 | 26.1 |
| Vtn | 0.0 | 0.0 | 0.0 | 0.0 | 0.0 | 0.0 | 0.9 |
| Vwa3a | 0.0 | 0.0 | 0.0 | 0.0 | 0.0 | 0.0 | 0.0 |
| Vwa5a | 26.6 | 18.0 | 14.7 | 16.6 | 14.1 | 11.3 | 38.3 |
| Vwa5b1 | 0.0 | 0.0 | 0.0 | 0.0 | 0.0 | 0.0 | 0.0 |
| Vwa5b2 | 0.0 | 0.1 | 0.1 | 0.1 | 0.1 | 0.3 | 0.1 |
| Vwa7 | 0.5 | 0.3 | 0.1 | 2.3 | 0.1 | 0.0 | 0.1 |
| Vwa9 | 9.7 | 8.1 | 6.3 | 8.3 | 8.8 | 11.7 | 8.2 |
| Vwc2 | 0.0 | 0.0 | 0.0 | 0.0 | 0.0 | 0.0 | 0.0 |
| Vwce | 0.0 | 0.0 | 0.0 | 0.0 | 0.0 | 0.1 | 0.1 |
| Vwde | 0.0 | 0.0 | 0.0 | 0.0 | 0.0 | 0.0 | 1.0 |
| Vwf | 0.1 | 0.0 | 0.1 | 0.0 | 0.1 | 0.2 | 0.1 |
| Wisp1 | 0.0 | 0.0 | 0.0 | 0.0 | 0.2 | 4.2 | 11.2 |
| Wisp2 | 0.1 | 0.0 | 0.1 | 0.2 | 0.1 | 1.7 | 46.9 |
| Wisp3 | 0.0 | 0.1 | 0.0 | 0.1 | 0.0 | 0.0 | 0.1 |
| Zp1 | 0.0 | 0.0 | 0.0 | 0.0 | 0.0 | 0.0 | 0.0 |
| Zp2 | 0.0 | 0.0 | 0.0 | 0.0 | 0.0 | 0.0 | 0.0 |
| Zp3 | 0.0 | 0.0 | 0.0 | 0.0 | 0.0 | 0.0 | 0.1 |
| Zp3r | 0.0 | 0.0 | 0.0 | 0.0 | 0.0 | 0.0 | 0.0 |
| Zpld1 | 0.0 | 0.0 | 0.0 | 0.0 | 0.0 | 0.0 | 0.0 |

**Supplementary Table 2.** Major cellular origin of basement membrane molecules

| Major cellular origin | Epidermis | Epidermis & dermis | Dermis |
|---|---|---|---|
| | Lama1 | Lamb1 | Lama2 |
| | Lama3 | Lamb2 | Lama4 |
| | Lama5 | Lamc1 | Lamc3 |
| | Lamb3 | Lamc2 | Col4a1 |
| | Col4a3 | Col7a1 | Col4a2 |
| | Col4a4 | Col8a2 | Col6a1 |
| | Col4a5 | Col18a1 | Col6a2 |
| | Col4a6 | Hspg2 | Col6a3 |
| | Col17a1 | Agrn | Col6a5 |
| | Egfl6 | Efemp1 | Col6a6 |
| | Emilin3 | Fbln1 | Col15a1 |
| | Fras1 | Hmcn1 | Nid1 |
| | Frem2 | Matn2 | Nid2 |
| | Npnt | Smoc1 | Colq |
| | Papln | Smoc2 | Egflam |
| | Tiagl1 | Sparc | Frem1 |
| | Vwa1 | Tgfbi | Ntn1 |
| | Vwa2 | Tnc | Ntn3 |
| | | | Ntn4 |
| | | | Ntn5 |
| | | | Ntng1 |
| | | | Thbs2 |
| | | | Thbs4 |

**Supplementary Table 3**. Summary of expression patterns of ECM genes in hair follicle epithelial compartments (shown in Fig. 3).

| Group | Basal | Lower isthmus | Upper bulge | Mid-bulge | Hair germ | ECM genes |
|---|---|---|---|---|---|---|
| 1 | + | + | + | + | + | *Lama5*, *Crim1*, *Lama3*, *Lamb2*, *Vwa5a*, *Creld2*, *Lamc1*, *Vwa9*, *Mfap3*, *Mfap1a*, *Ntn4* |
| 2 | + | + | + | + | - | *Gas6*, *Col4a5*, *Creld1* |
| 3 | - | + | + | + | + | *Col20a1* |
| 4 | + | - | + | + | + | *Ltbp4*, *Lamc2*, *Frem2* |
| 5 | + | + | + | - | - | *Smoc2* |
| 6 | - | + | + | + | - | *Igfbp4*, *Aspn* |
| 7 | - | - | + | + | + | *Agrn*, *Fbln1*, *Ddx26b*, *Hspg2*, *Col4a1*, *Col4a2*, *Ltbp1*, *Col5a2*, *Efemp2*, *Col1a2*, *Aebp1*, *Tsku* |
| 8 | + | - | - | + | + | *Cyr61*, *Col16a1*, *Adamtsl5* |
| 9 | + | + | - | + | - | *Igfbp3*, *Thbs2*, *Col4a6* |
| 10 | + | - | + | + | - | *Lamb3* |
| 11 | - | + | - | + | + | *Slit3*, *Spock2* |
| 12 | + | - | + | - | + | *Efemp1* |
| 13 | - | + | + | - | - | *Lama4* |
| 14 | - | - | + | - | + | *Abi3bp*, *Mgp*, *Fn1*, *Mfap2*, *Lamb1*, *Bgn*, *Pxdn*, *Col1a1*, *Fras1* |
| 15 | - | - | + | + | - | *Vwa2*, *Igfbp7*, *Npnt*, *Egfl6*, *Crispld1*, *Emid1*, *Col5a1* |
| 16 | + | - | - | + | - | *Col17a1*, *Tgfbi*, *Col7a1* |
| 17 | - | - | - | + | + | *Tnc*, *Hmcn1*, *Thbs3*, *Fbln2* |
| 18 | + | - | - | - | - | *Tnxb*, *Col23a1*, *Rspo3*, *Ntng2*, *Lama1* |
| 19 | - | + | - | - | - | *Igfbp2* |
| 20 | - | - | + | - | - | *Col4a4*, *Col4a3*, *Fmod* |
| 21 | - | - | - | + | - | *Postn*, *Ctgf*, *Ecm1*, *Sparc*, *Col18a1*, *Ltbp3*, *Col6a1*, *Adamtsl4*, *Col8a2*, *Ltbp2*, *Mfge8*, *Col6a2*, *Vwa1*, *Igfbp5*, *Igfbp6*, *Matn2*, *Spon2*, *Papln*, *Col12a1*, *Pcolce2*, *Smoc1*, *Vit*, *Emilin3*, *Chadl* |
| 22 | - | - | - | - | + | *Thbs1*, *Tnfaip6*, *Omd*, *Crispld2*, *Spp1*, *Col6a3*, *Tinagl1*, *Dcn*, *Cilp*, *Spon1*, *Thsd4*, *Igfals*, *Podn*, *Sned1*, *Pcolce*, *Dpt*, *Igsf10*, *Fgl2*, *Col3a1*, *Lamc3*, *Rspo1*, *Srgn*, *Nov*, *Fgg*, *Lama2* |

**Supplementary Table 4.** List of ECM antibodies screened for reactivity

**Common ECMs**

| Gene symbol (mean FPKM in epidermis) | Protein name (Abbreviation) | Product ID or clone name of tested antibody | Manufacturer | ECM-like localization | Tissue localization around hair follicle epithelium |
|---|---|---|---|---|---|
| Lama5 (45.9) | laminin subunit alpha-5 (laminin α5) | CUK-1185-006 | Fujiwara Lab (in house) | Yes | Entire BM zone (weak at mid-bulge) |
| Crim1 (40.3) | cysteine-rich motor neuron 1 protein (CRIM-1) | HPA000556 | Atlas Antibodies | No | |
| | | AF1917 | R&D | No (nuclei) | |
| | | GTX51530 | GeneTex | No | |
| | | ExCrim1 | Fujiwara Lab (in house) | Yes | BM zone of mid-bulge |
| Lama3 (31.5) | laminin subunit alpha-3 (laminin α3) | ab14509 (as laminin 332) | abcam | Yes | Entire BM zone |
| | | LS-C670558 | LSBio | No | |
| Lamb2 (21.6) | laminin subunit beta-2 (laminin β2) | ab21096 | abcam | No | |
| | | B24-N8-D6 | Sekiguchi Lab (Manabe et al., PNAS, 2008) | Yes | Entire BM zone |
| Vwa5a (18.0) | von Willebrand factor A domain-containing protein 5A | Not tested | | | |
| Creld2 (17.3) | protein disulfide isomerase Creld2 | Not tested | | | |
| Lamc1 (12.3) | laminin subunit gamma-1 (laminin γ1) | MAB1914P (A5) | Merck | Yes | Entire BM zone |
| Ints14 (Vwa9) (8.2) | integrator complex subunit 14 | Not tested | | | |
| Mfap3 (5.3) | microfibril-associated glycoprotein 3 | Not tested | | | |
| Mfap1a (4.6) | microfibrillar-associated protein 1A | Not tested | | | |
| Ntn4 (3.8) | netrin-4 | AF1132 | R&D | Yes | Entire BM zone (weak at mid-bulge) |

**ECMs preferentially expressed in the upper bulge region**

| Gene symbol (mean | Protein name (Abbreviation) | Product ID or clone name of | Manufacturer | ECM-like localization | Tissue localization around hair follicle epithelium |
|---|---|---|---|---|---|

| | | | | | |
|---|---|---|---|---|---|
| **FPKM in epidermis)** | | **tested antibody** | | | |
| Col4a4 (2.5) | collagen alpha-3 (IV) chain | RH42 | Shigei Med. Res. Inst. | Yes | BM zone of upper bulge |
| Col4a3 (2.2) | collagen alpha-3 (IV) chain | H31 | Shigei Med. Res. Inst. | Yes | BM zone of upper bulge |
| Fmod (1.9) | fibromodulin | Not tested | | | |

**ECMs preferentially expressed in the mid-bulge region**

| Gene symbol (mean FPKM in epidermis) | Protein name (Abbreviation) | Product ID or clone name of tested antibody | Manufacturer | ECM-like localization | Tissue localization around hair follicle epithelium |
|---|---|---|---|---|---|
| Postn (551.5) | periostin | ab14041 | abcam | Yes | BM zone of mid-bulge |
| Ctgf (248.5) | CCN family member 2 (CTGF) | ab6992 | abcam | Yes | Upper bulge lanceolate complex |
| Ecm1 (184.5) | extracellular matrix protein 1 | 11521-1-AP | Proteintech | No | |
| | | AF4428 | R&D | No | |
| | | AF3937 | R&D | No | |
| Sparc (71.4) | SPARC | Ab14174 | abcam | No | |
| | | MAB942-SP | R&D | No | |
| | | AF941 | R&D | No (nuclei) | |
| Col18a1 (58.4) | collagen alpha-1(XVIII) chain | AF1098 | R&D | Yes | Entire BM zone (strong at mid-bulge) |
| Ltbp3 (50.9) | latent-transforming growth factor beta-binding protein 3 (LTBP-3) | ABT316 | Merck | Yes | BM zone of mid-bulge |
| | | sc-390913 AF488 | Santa Cruz | Yes | BM zone of mid-bulge |
| Col6a1 (43.3) | collagen alpha-1(VI) chain | Ab6588 | abcam | Yes | Around mid-bulge, HG and DP |
| | | HPA029401 | Sigma | No | |
| Adamtsl4 (31.5) | ADAMTS-like protein 4 (adamtsl4) | Not tested | | | |
| Col8a2 (28.4) | collagen alpha-2(VIII) chain | 034099 | USBiological | No | |
| Ltbp2 (27.6) | latent-transforming growth factor beta-binding protein 2 (LTBP-2) | sc-48759 | Santa Cruz | Yes | BM zone of mid-bulge and HG regions |
| Mfge8 (18.7) | lactadherin | 18A2-G10 | MBL | No | |

| | | | | | |
|---|---|---|---|---|---|
| Col6a2 (17.7) | collagen alpha-2(VI) chain | sc-8360 | Santa Cruz | No | |
| Vwa1 (12.1) | von Willebrand factor A domain-containing protein 1 | | Sekiguchi Lab (Manabe et al., PNAS, 2008) | Yes | Entire BM zone (strong at mid-bulge) |
| Igfbp5 (10.9) | insulin-like growth factor-binding protein 5 (IGFBP-5) | AF578 | R&D | Yes | Upper bulge sensory neuron |
| Igfbp6 (10.4) | insulin-like growth factor-binding protein 5 (IGFBP-6) | Sc-13094 | Santa Cruz | No | |
| Matn2 (9.1) | matrilin-2 | AF3044 | R&D | Yes | BM zones of isthmus, upper bulge and mid-bulge regions |
| Spon2 (8.1) | spondin-2 | 20513-1-AP | Proteintech | No | |
| | | PA006509 | Cusabio | No | |
| Papln (7.4) | papilin | | Sekiguchi Lab (Manabe et al., PNAS, 2008) | Yes | Entire BM zone (strong at mid-bulge) |
| Col12a1 (6.5) | collagen alpha-1(XII) chain | STJ92375 | St John's | No | |
| Pcolce2 (3.1) | procollagen C-endopeptidase enhancer 2 | 10607-1-AP | Proteintech | No | |
| Smoc1 (2.1) | SPARC-related modular calcium-binding protein 1 (SMOC-1) | AF5550 | R&D | Yes | BM zone at HG-DP interface |
| Vit (2.0) | vitrin | NBP2-49453 | Novus | No | |
| Emilin3 (1.7) | EMILIN-3 | Not tested | | | |
| Chadl (1.5) | chondroadherin-like protein | Not tested | | | |

**ECMs expressed in the mid-bulge region and basal cell**

| **Gene symbol (mean FPKM in epidermis)** | **Protein name (Abbreviation)** | **Product ID or clone name of tested antibody** | **Manufacturer** | **ECM-like localization** | **Tissue localization around hair follicle epithelium** |
|---|---|---|---|---|---|
| Col17a1 (934.2) | collagen alpha-1(XVII) chain | R311 | Hirako Lab (Hirako et al., J. Biochem., 2003) | Yes | BM zone of mid-bulge |
| Tgfbi (262.1) | transforming growth factor- | | Sekiguchi Lab (Manabe et al., | Yes | Entire BM zone |

| | | | | | |
|---|---|---|---|---|---|
| | beta-induced protein ig-h3 | | PNAS, 2008) | | |
| Col7a1 (58.9) | collagen alpha-1(VII) chain | LS-B10205 | LSBio | No | |
| | | 234192 | Merck | Yes | Entire BM zone |

**ECMs preferentially expressed in the hair germ region**

| Gene symbol (mean FPKM in epidermis) | Protein name (Abbreviation) | Product ID or clone name of tested antibody | Manufacturer | ECM-like localization | Tissue localization around hair follicle epithelium |
|---|---|---|---|---|---|
| Thbs1 (381.9) | thrombospondin-1 | sc-59887 | Santa Cruz | No | |
| Tnfaip6 (12.0) | tumor necrosis factor-inducible gene 6 protein | HPA050884 | Sigma | No | |
| | | PA5-62259 | Thermo Fisher | No | |
| Omd (4.0) | osteomodulin | Not tested | | | |
| Crispld2 (3.3) | cysteine-rich secretory protein LCCL domain-containing 2 | Not tested | | | |
| Spp1 (3.2) | osteopontin | ab8448 | abcam | No | |
| Col6a3 (3.1) | collagen alpha-3(VI) chain | PAB17517 | Abnova | No | |
| | | HPA010080 | Sigma | Yes | Lateral side BM zone of HG |
| Tinagl1 (2.6) | tubulointerstitial nephritis antigen-like | Not tested | | | |
| Dcn (2.5) | decorin | AF1060 | R&D | Yes | No (interstitial fibrils) |
| Spon1 (2.3) | spondin-1 | ab40797 | abcam | No | |
| | | AF3135 | R&D | Yes | BM zone at HG-DP interface |
| Cilp (2.3) | cartilage intermediate layer protein 1 | Not tested | | | |
| Thsd4 (2.1) | thrombospondin type-1 domain-containing protein 4 (ADAMTS-like protein 6) | | Sekiguchi Lab (Manabe et al., PNAS, 2008) | Yes | BM zone at HG-DP interface |
| Podn (2.0) | podocan | Not tested | | | |
| Igfals (2.0) | insulin-like growth factor-binding protein complex acid labile subunit | Not tested | | | |
| Dpt | dermatopontin | LS-C167524 | LSBio | No | |

| | | | | | |
|---|---|---|---|---|---|
| (1.8) | | PAC432Hu01 | Cloud-Clone | No | |
| Sned1 (1.8) | sushi, nidogen and EGF-like domain-containing protein 1 | HPA036415 | Atlas Antibodies | No | |
| Pcolce (1.8) | procollagen C-endopeptidase enhancer 1 | Not tested | | | |
| Igsf10 (1.7) | immunoglobulin superfamily member 10 | Not tested | | | |
| Fgl2 (1.5) | fibroleukin | HPA021011 | Sigma | No | |
| Col3a1 (1.5) | collagen alpha-1(III) chain | ab7778 | abcam | Yes | Broad BM zone except mid-bulge |
| Lamc3 (1.3) | laminin subunit gamma-3 (laminin γ3) | sc-25719 | Santa Cruz | No | |
| | | C38-N4-F4 | Sekiguchi Lab (Manabe et al., PNAS, 2008) | Yes | BM zone at HG-DP interface |
| Rspo1 (1.3) | R-spondin-1 | sc-49092-R | Santa Cruz | No | |
| | | HPA046154 | Atlas Antibodies | No | |
| | | AF3474 | R&D | No | |
| Srgn (1.2) | serglycin | C-11 | Santa Cruz | No | |
| | | H00005552-M03 | Abnova | No | |
| Ccn3 (Nov) (0.9) | CCN family member 3 (nephroblastoma overexpressed) | Not tested | | | |
| Lama2 (0.7) | laminin subunit alpha-2 (laminin α2) | 4H8-2 | abcam | Yes | BM zone of HG and peripheral nerve |
| Fgg (0.7) | fibrinogen gamma chain | Not tested | | | |

**ECMs expressed in the hair germ and upper bulge regions**

| **Gene symbol (mean FPKM in epidermis)** | **Protein name (Abbreviation)** | **Product ID or clone name of tested antibody** | **Manufacturer** | **ECM-like localization** | **Tissue localization around hair follicle epithelium** |
|---|---|---|---|---|---|
| Abi3bp (45.1) | target of Nesh-SH3 | | Sekiguchi Lab (Manabe et al., PNAS, 2008) | No | |
| Mgp (21.3) | matrix Gla protein | Not tested | | | |
| Fn1 (17.2) | fibronectin | ab23750 | abcam | Yes | Around mid-bulge, HG and DP |

| | | | | | |
|---|---|---|---|---|---|
| Mfap2 (13.4) | microfibrillar-associated protein 2 (MFAP-2) | Not tested | | | |
| Lamb1 (11.6) | laminin subunit beta-1 (laminin β1) | LT3 | GeneTex | Yes | BM zone at HG-DP interface and peripheral nerve |
| Bgn (10.8) | biglycan | AF2667 | R&D | Yes | Peripheral nerve |
| Pxdn (4.6) | peroxidasin homolog | sc-168598 | Santa Cruz | Yes | Peripheral nerve |
| Col1a1 (2.8) | collagen alpha-1(I) chain | Not tested | | | |
| Fras1 (2.2) | extracellular matrix protein FRAS1 | Sc-98444 | Santa Cruz | Yes | Peripheral nerve |

**ECMs preferentially expressed in the dermal papilla**

| Gene symbol (FPKM in the dermal papilla) | Protein name (Abbreviation) | Product ID or clone name of tested antibody | Manufacturer | ECM-like localization | Tissue localization around the dermal papilla |
|---|---|---|---|---|---|
| Igfbp3 (1667.3) | insulin-like growth factor-binding protein 3 (IGFBP-3) | Not tested | | | |
| Spp1 (645.2) | osteopontin | ab8448 | abcam | No | |
| Spon1 (223.2) | spondin-1 | ab40797 | abcam | No | |
| | | AF3135 | R&D | Yes | Interface BM zone, hook and mesh BMs |
| Col23a1 (111.8) | collagen alpha-1(XXIII) chain | NBP2-38111 | Novus | No | |
| Lamc3 (84.6) | laminin subunit gamma-3 (laminin γ3) | sc-25719 | Santa Cruz | No | |
| | | C38-N4-F4 | Sekiguchi Lab (Manabe et al., PNAS, 2008) | Yes | BM zone at HG-DP interface |
| Srgn (76.9) | serglycin | C-11 | Santa Cruz | No | |
| | | H00005552-M03 | Abnova | No | |
| Tnn (57.6) | tenascin-N | ab121887 | abcam | Yes | Interface BM zone, hook and mesh BMs |
| Ints6l (Ddx26b) (52.4) | integrator complex subunit 6-like (DDX26B) | Not tested | | | |
| Spock1 (41.7) | testican-1 | 12512-1-AP | Proteintech | No | |
| Col6a6 (41.0) | collagen alpha-6(VI) chain | NBP2-14546 | Novus | No | |
| | | HPA045239 | Sigma | Yes | Interface BM zone (and isthmus BM |

| | | | | | |
|---|---|---|---|---|---|
| | | | | | zone) |
| Thsd4 (38.6) | thrombospondin type-1 domain-containing protein 4 (ADAMTS-like protein 6) | | Sekiguchi Lab (Manabe et al., PNAS, 2008) | Yes | Hook BM |
| Col13a1 (38.2) | collagen alpha-1(XIII) chain | HPA050392 | Atlas Antibodies | Yes | Interface BM zone and hook BM |
| | | AF4627 | R&D | Yes | Interface BM zone and hook BM |
| Ndnf (18.8) | protein NDNF | AP5812a | Abgent | No | |
| | | sc-242196 | Santa Cruz | No | |
| | | LS-C319946 | LSBio | No | |
| Crispld1 (17.0) | cysteine-rich secretory protein LCCL domain-containing 1 | ab123039 | abcam | No | |
| Rspo3 (15.2) | R-spondin-3 | 17193-1-AP | Proteintech | Yes | Around DP |
| Col6a5 (11.3) | collagen alpha-5(VI) chain | HPA043138 | Atlas Antibodies | Yes | No (peripheral nerve) |
| Col27a1 (10.7) | collagen alpha-1(XXVII) chain | ab179753 | abcam | No | |
| | | PA5-54535 | Thermo Fisher | No | |
| Lamc2 (10.6) | laminin subunit gamma-2 (laminin γ2) | Not tested | | | |
| Smoc1 (5.4) | SPARC-related modular calcium-binding protein 1 (SMOC-1) | AF5550 | R&D | Yes | Interface BM zone |
| Edil3 (5.4) | EGF-like repeat and discoidin I-like domain-containing protein 3 | ab198003 | abcam | No | |
| Rspo2 (3.3) | R-spondin-2 | 879712R | R&D | No | |
| Matn1 (3.1) | cartilage matrix protein (matrilin 1) | HPA028580 | Atlas Antibodies | No | |

**Integrin proteins expressed in the hair follicle**

| **Gene symbol** | **Protein name** | **Product ID or clone name of tested antibody** | **Manufacturer** | **Cell membrane localization** | **Tissue localization in the hair follicle** |
|---|---|---|---|---|---|
| Itgb1 | integrin β1 | HMb1-1 | BioLegend | Yes | Hair germ cells, cells around hook BM and peripheral nerve |

| | | | | | |
|---|---|---|---|---|---|
| Itgav | integrin αv | AF1219 | R&D | Yes | Cells around hook BM |
| Itga6 | integrin α6 | GoH3 | BioLegend | Yes | All basal cells and cells around hook BM |
| Itgb4 | integrin β4 | HPA036348 | Sigma | Yes | Interfollicular epidermis, isthmus and hair germ cells and peripheral nerve |
| Itga3 | integrin α3 | AF2787 | R&D | Yes | Isthmus and hair germ cells and peripheral nerve |
| Itga5 | integrin α5 | ab221606 | abcam | Yes | Interfollicular epidermis and hair germ cells |
| Itga8 | integrin α8 | AF4076 | R&D | Yes | Interface zone between hair germ and dermal papilla |
| Itga9 | integrin α9 | AF3827 | R&D | Yes | Cells around mesh BM |
| Itgb8 | integrin β8 | sc-25714 | Sanat Cruz | No | |

**Supplementary Table 5.** Primer sequences used for qRT-PCR

| Target genes | Primer sequence (forward) | Primer sequence (reverse) |
|---|---|---|
| *Gapdh* | TGCCAGCCTCGTCCCGTAG | CGGCCTTGACTGTGCCGTTG |
| *Itga6* | CGGGCACTCAGGTTCGAGTGA | AGCCTTGTGATAGGTGGCATCGT |
| *Lrig1* | TGTGTGTCTGAGAGACCCGAGC | CAGAGCCACTGTGTGCTGTTGT |
| *Lgr6* | GGCTCCGAATCCTGGAGCTGT | CTCAGGGTGGATGGCACGGA |
| *Gli1* | ATCTCCGGGCGGTTCCTACG | TGACTTCAGCTGGCAGGTTGC |
| *Bdnf* | GCGGCGCCCATGAAAGAAGT | GGCCGAACCTTCTGGTCCTCA |
| *Cd34* | TGGCGCTGGGTAGCTCTCTG | TAGTCTCTGAGATGGCTGGTGTGG |
| *Cdh3* | TGGGGAAAGTAGCCTTGGCTGG | CAGCCTCTGAGGGAAGGGACC |
| *Lef1* | GCCGGGATGCCCCAACTTTC | GACCACCTCATGCCCGTTGC |
| *Pdgfra* | TGCGGGTTTTGAGCCCATTACT | CCGGCCCTGTGAGGAGACAG |
| *Itga8* | TCTTGTGCAGTGGGTCGCCT | CCGACGTCTTAACCGCTGTGC |
| *eGFP* | TGGTGCCCATCCTGGTCGAC | GAAGCACTGCACGCCGTAGG |

**Supplementary Table 6.** RNA-seq read and mapping statistics

| | Raw Read Count | Trimmed Read Count | rDNA Read Count | Multi-Mapped Read Count | Uniquely mapped Read Count |
|---|---|---|---|---|---|
| Mouse_Basal_ Cells_Rep1 | 11,757,494 | 11,278,061 | 148,700 | 170,077 | 10,456,332 |
| Mouse_Basal_ Cells_Rep2 | 11,245,045 | 10,668,752 | 73,729 | 134,777 | 10,051,376 |
| Mouse_Basal_ Cells_Rep3 | 11,313,239 | 10,747,103 | 108,596 | 146,602 | 10,089,518 |
| Mouse_Bulge_ StemCells_Rep1 | 11,735,034 | 11,181,377 | 64,614 | 183,557 | 10,520,864 |
| Mouse_Bulge_ StemCells_Rep2 | 11,917,049 | 11,515,851 | 54,154 | 175,176 | 10,889,932 |
| Mouse_Bulge_ StemCells_Rep3 | 13,417,100 | 12,895,113 | 88,655 | 218,495 | 12,090,872 |
| Mouse_Cdh3+_ Cells_Rep1 | 10,875,593 | 10,409,342 | 68,193 | 177,407 | 9,574,901 |
| Mouse_Cdh3+_ Cells_Rep2 | 11,565,616 | 10,993,093 | 65,001 | 185,981 | 10,079,620 |
| Mouse_Cdh3+_ Cells_Rep3 | 11,405,265 | 10,746,768 | 76,143 | 206,384 | 9,790,096 |
| Mouse_Cdh3+_ Cells_Rep4 | 11,759,881 | 11,186,543 | 85,902 | 191,154 | 10,274,371 |
| Mouse_Gli1+_ Cells_Rep1 | 12,063,488 | 11,478,537 | 57,630 | 178,646 | 10,751,521 |
| Mouse_Gli1+_ Cells_Rep2 | 19,762,234 | 18,804,713 | 178,685 | 303,848 | 17,446,776 |
| Mouse_Gli1+_ Cells_Rep3 | 19,543,117 | 18,568,695 | 122,017 | 287,214 | 17,378,904 |
| Mouse_Lgr6+_ Cells_Rep1 | 20,044,455 | 19,058,807 | 121,507 | 307,804 | 17,798,594 |
| Mouse_Lgr6+_ Cells_Rep2 | 19,756,354 | 18,736,071 | 197,458 | 298,652 | 17,386,391 |
| Mouse_Lgr6+_ Cells_Rep3 | 19,639,299 | 18,680,969 | 217,502 | 298,146 | 17,210,635 |
| Mouse_Lef1+_ Cells_Rep1 | 13,353,072 | 12,447,662 | 90,361 | 163,390 | 11,573,835 |
| Mouse_Lef1+_ Cells_Rep2 | 11,534,943 | 10,876,371 | 69,732 | 150,330 | 10,020,573 |
| Mouse_Lef1+_ Cells_Rep3 | 11,479359 | 10,679,409 | 55,674 | 123,456 | 10,001,747 |
| Mouse_Lef1+_ Cells_Rep4 | 10,991,615 | 10,405,782 | 42,283 | 117,783 | 9,796,296 |
| Mouse_Pdgfa+_ Cells_Rep1 | 21,043,519 | 20,002,460 | 197,994 | 308,101 | 18,559,766 |
| Mouse_Pdgfa+_ Cells_Rep2 | 20,727,828 | 19,678415 | 211,148 | 324,166 | 18,190,226 |
| Mouse_Pdgfa+_ Cells_Rep3 | 20,638,004 | 19,549,144 | 214,785 | 339,207 | 17,998,448 |

**Supplementary Table 7.** Antibodies used in this study, their specific dilutions and tissue fixation methods

| Designation | Host species | Source or reference | Identifiers | Use and dilution | Fixation |
|---|---|---|---|---|---|
| **FACS** | | | | | |
| CD45-PE-Cy7 | Rat | eBioscience | 30-F11 | FACS (1:100) | no |
| TER-119-PE-Cy7 | Rat | eBioscience | TER119 | FACS (1:100) | no |
| CD31-PE-Cy7 | Rat | eBioscience | 390 | FACS (1:100) | no |
| Sca-1-PerCP-Cy5.5 | Rat | eBioscience | D7 | FACS (1:100) | no |
| CD34-eFluor660 | Rat | eBioscience | RAM34 | FACS (1:100) | no |
| CD49f-PE | Rat | eBioscience | GoH3 | FACS (1:100) | no |
| **ECM-related** | | | | | |
| ABI3BP | Rabbit | Sekiguchi lab. (Manabe et al., PNAS, 2008) | serum | IHC (1:8) | PFA |
| adamtsl-6 | Rabbit | Sekiguchi lab. (Manabe et al., PNAS, 2008) | serum | IHC (1:500) | PFA |
| biglycan | Goat | R&D | AF2667 | IHC (10 µg/ml) | PFA |
| CD34 | Rat | Hycult Biotech | HM1015 (MEC14.7) | IHC (2 µg/ml) | PFA/acetone +acid |
| CD34 | Rabbit | abcam | ab81289 | IHC (10.3 µg/ml) | PFA/acetone +acid |
| collagen III | Rabbit | abcam | ab7778 | IHC (50 µg/ml) | PFA |
| COL4A1 | Rat | Shigei Med. Res. Inst. | H11 | IHC (1:20) | Acetone+acid |
| COL4A3 | Rat | Shigei Med. Res. Inst. | H31 | IHC (1:10) | Acetone+acid |
| COL4A4 | Rat | Shigei Med. Res. Inst. | RH42 | IHC (1:10) | Acetone+acid |
| collagen VI | Rabbit | abcam | Ab6588 | IHC (50 µg/ml) | PFA |
| COL6A1 | Rabbit | Sigma | HPA029401 | IHC (2.5 µg/ml) | PFA |
| COL6A2 | Rabbit | Santa Cruz | sc-8360 | IHC (1 µg/ml) | PFA |
| COL6A3 | Rabbit | Abnova | PAB17517 | IHC (10 µg/ml) | PFA |
| COL6A3 | Rabbit | Sigma | HPA010080 | IHC (1.25 µg/ml) | PFA |
| COL6A5 | Rabbit | Atlas Antibodies | HPA043138 | IHC (0.5 µg/ml) | PFA |
| COL6A6 | Rabbit | Novus | NBP2-14546 | IHC (30 µg/ml) | PFA |
| COL6A6 | Rabbit | Sigma | HPA045239 | IHC (1 µg/ml) | PFA |
| COL7A1 | Rabbit | LSBio | LS-B10205 | IHC (20 µg/ml) | PFA |
| COL7A1 | Rabbit | Merck | 234192 | IHC (1:40) | PFA |
| COL8A2 | Rabbit | USBiological | 034099 | IHC (2.3 or 11.4 µg/ml) | PFA/acetone +acid |
| COL12A1 | Rabbit | St John's | STJ92375 | IHC (12.5 µg/ml) | PFA |

| | | | | | |
|---|---|---|---|---|---|
| COL13A1 | Rabbit | Atlas Antibodies | HPA050392 | IHC (2.5 μg/ml) | PFA |
| COL13A1 | Sheep | R&D | AF4627 | IHC (5.6 μg/ml) | PFA |
| COL17A1 | Mouse | Hirako Lab (Hirako et al., J. Biochem., 2003) | R311 | IHC (1:1) | PFA |
| COL18A1 | Goat | R&D | AF1098 | IHC (4 μg/ml) | PFA |
| COL23A1 | Rabbit | Novus | NBP2-38111 | IHC (2.5 μg/ml) | PFA/acetone +acid |
| COL27A1 | Rabbit | abcam | Ab179753 | IHC (25 μg/ml) | PFA |
| COL27A1 | Rabbit | Thermo Fisher | PA5-54535 | IHC (2.5 μg/ml) | PFA |
| CRIM1 | Rabbit | Atlas Antibodies | HPA000556 | IHC (5 or 2 μg/ml) | PFA/acetone +acid |
| CRIM1 | Goat | R&D | AF1917 | IHC (2 μg/ml) | PFA/acetone +acid |
| CRIM1 | Rabbit | GeneTex | GTX51530 | IHC (10 μg/ml) | PFA |
| CRIM1 | Rabbit | Fujiwara Lab. (this paper) | serum | IHC (1:400) | PFA |
| CRISPLD1 | Rabbit | abcam | ab123039 | IHC (25 μg/ml) | PFA |
| CTGF | Rabbit | abcam | ab6992 | IHC (25 μg/ml) | PFA |
| CTGF | Goat | R&D | AF660 | IHC (12.6 μg/ml) | PFA |
| decorin | Goat | R&D | AF1060 | IHC (2 μg/ml) | PFA |
| dermatopontin | Rabbit | LSBio | LS-C167524 | IHC (1:20) | PFA |
| dermatopontin | Rabbit | Cloud-Clone | PAC432Hu01 | IHC (20 μg/ml) | PFA |
| ECM1 | Rabbit | Proteintech | 11521-AP | IHC (9.3 or 1.9 μg/ml) | PFA/acetone +acid |
| ECM1 | Goat | R&D | AF4428 | IHC (0.4 or 4 μg/ml) | PFA/acetone +acid |
| ECM1 | Sheep | R&D | AF3937 | IHC (2 μg/ml) | PFA |
| EDIL3 | Rabbit | abcam | Ab198003 | IHC (1:100) | PFA |
| EDIL3 | Rabbit | Proteintech | 12580-1-AP | IHC (6.7 μg/ml) | PFA |
| FGL2 | Rabbit | Sigma | HPA021011 | IHC (2.5 μg/ml) | PFA |
| fibronectin | Rabbit | abcam | ab23750 | IHC (4 μg/ml) | PFA |
| fibulin-2 | Rabbit | Atlas Antibodies | HPA001934 | IHC (10 μg/ml) | PFA/acetone +acid |
| FRAS1 | Rabbit | Santa Cruz | sc-98444 | IHC (10 or 2 μg/ml) | PFA/acetone +acid |
| IGFBP5 | Goat | R&D | AF578 | IHC (1 μg/ml) | PFA |
| IGFBP6 | Rabbit | Santa Cruz | sc-13094 | IHC (5 μg/ml) | PFA |
| integrin α3 | Goat | R&D | AF2787 | IHC (2 μg/ml) | PFA |
| integrin α5 | Rabbit | abcam | Ab221606 | IHC (2.1 μg/ml) | PFA |
| integrin α6 | Rat | Biolegend | GoH3 | IHC (10 μg/ml) | PFA |
| integrin α8 | Goat | R&D | AF4076 | IHC (0.8 μg/ml) | PFA |
| integrin α9 | Goat | R&D | AF3827 | IHC (2 μg/ml) | PFA |
| integrin αv | Goat | R&D | AF1219 | IHC (10 μg/ml) | PFA |
| integrin β1 | Hamster | Biolegend | HMb1-1 | IHC (10 μg/ml) | PFA |
| integrin β4 | Rabbit | Sigma | HPA036348 | IHC (2 μg/ml) | PFA |
| integrin β8 | Rabbit | Santa Cruz | sc-25714 | IHC (5 μg/ml) | PFA |
| laminin-332 | Rabbit | abcam | Ab14509 | IHC (5 μg/ml) | PFA |
| laminin α2 | Rat | abcam | 4H8-2 | IHC (20.8 | PFA |

| | | | | | |
|---|---|---|---|---|---|
| | | | | μg/ml) | |
| laminin α3 | Rabbit | LSBio | LS-C670558 | IHC (8.1 μg/ml) | PFA |
| laminin α4 | Goat | R&D | AF3837 | IHC (5 μg/ml) | PFA |
| laminin α5 | Rabbit | Fujiwara Lab. (this paper) | CUK-1185-006 | IHC (0.09 μg/ml) | PFA |
| laminin β1 | Rat | GeneTex | LT3 | IHC (0.7 μg/ml) | PFA |
| laminin β2 | Mouse | abcam | Ab210956 | IHC (1:10) | PFA |
| laminin β2 | Rat | Sekiguchi lab. (Manabe et al., PNAS, 2008) | B24-N8-D6 | IHC (3.7 μg/ml) | Acetone+acid |
| laminin γ1 | Rat | Merck | MAB1914P | IHC (1 μg/ml) | PFA |
| laminin γ3 | Rabbit | Santa Cruz | sc-25719 | IHC (2 or 4 μg/ml) | PFA/acetone +acid |
| laminin γ3 | Rat | Sekiguchi lab. (Manabe et al., PNAS, 2008) | C38-N4-F4 | IHC (3.3 μg/ml) | Acetone+acid |
| LTBP2 | Rabbit | Santa Cruz | sc-48759 | IHC (5 μg/ml) | PFA |
| LTBP3 | Rabbit | Merck | ABT316 | IHC (12.5 μg/ml) | PFA |
| LTBP3 | Mouse | Santa Cruz | sc-390913 AF488 | IHC (10 μg/ml) | PFA |
| matrilin-1 | Rabbit | Atlas Antibodies | HPA028580 | IHC (2.5 μg/ml) | PFA |
| matrilin-2 | Goat | R&D | AF3044 | IHC (25.1 μg/ml) | PFA |
| MFGE8 | Hamster | MBL | 18A2-G10 | IHC (1 μg/ml) | PFA |
| NDNF | Rabbit | Abgent | AP5812a | IHC (25 μg/ml) | PFA |
| NDNF | Goat | Santa Cruz | sc-242196 | IHC (5 μg/ml) | PFA |
| NDNF | Rabbit | LSBio | LS-C319946 | IHC (6.4 μg/ml) | PFA |
| netrin-4 | Goat | R&D | AF1132 | IHC (5 μg/ml) | PFA |
| nidogen-1 | Rat | Merck | MAB1886 | IHC (4 μg/ml) | PFA |
| nidogen-2 | Goat | R&D | AF3385 | IHC (0.8 μg/ml) | PFA |
| osteopontin | Rabbit | abcam | ab8448 | IHC (1:500) | PFA |
| papilin | Rabbit | Sekiguchi lab. (Manabe et al., PNAS, 2008) | serum | IHC (1:40) | PFA |
| PCOLCE2 | Rabbit | Proteintech | 10607-1-AP | IHC (12.3 μg/ml) | PFA |
| periostin | Rabbit | abcam | ab14041 | IHC (12.5 μg/ml) | PFA |
| perlecan | Rat | Merck | MAB1948P | IHC (1:500) | PFA |
| peroxidasin | Goat | Santa Cruz | sc-168598 | IHC (0.2 μg/ml) | PFA |
| plectin | Guinea pig | Progen | GP21 | IHC (1:1,000) | PFA |
| R-spondin-1 | Rabbit | Santa Cruz | sc-49092-R | IHC (10 μg/ml) | PFA |
| R-spondin-1 | Rabbit | Atlas Antibodies | HPA046154 | IHC (20 μg/ml) | PFA |
| R-spondin-1 | Goat | R&D | AF3474 | IHC (1 or 2 μg/ml) | PFA/acetone +acid |
| R-spondin-2 | Rat | R&D | 879712R | IHC (2.5 μg/ml) | PFA |
| R-spondin-3 | Rabbit | Proteintech | 17193-1-AP | IHC (9.3 or 3.7 μg/ml) | PFA/acetone +acid |
| serglycin | Mouse | Santa Cruz | C-11 | IHC (5 μg/ml) | PFA |

| | | | | | |
|---|---|---|---|---|---|
| serglycin | Mouse | Abnova | H00005552-M03 | IHC (12 μg/ml) | PFA |
| SMOC1 | Goat | R&D | AF5550 | IHC (25 μg/ml) | PFA |
| SNED1 | Rabbit | Atlas Antibodies | HPA036415 | IHC (2.5 μg/ml) | PFA |
| SPARC | Rabbit | abcam | Ab14174 | IHC (1:200) | PFA |
| SPARC | Rat | R&D | MAB942-SP | IHC (25 μg/ml) | PFA |
| SPARC | Goat | R&D | AF941 | IHC (5 μg/ml) | PFA |
| SPOCK1 | Rabbit | Proteintech | 12512-1-AP | IHC (11 μg/ml) | PFA |
| spondin-1 | Rabbit | abcam | ab40797 | IHC (2 μg/ml) | PFA |
| spondin-1 | Goat | R&D | AF3135 | IHC (0.4 μg/ml) | PFA |
| spondin-2 | Rabbit | Proteintech | 20513-1-AP | IHC (4.1or 20.7 μg/ml) | PFA/acetone +acid |
| spondin-2 | Rabbit | Cusabio | PA006509 | IHC (25 μg/ml) | PFA |
| tenascin-N | Rabbit | abcam | ab121887 | IHC (5 μg/ml) | PFA |
| TGFBI | Rabbit | Sekiguchi lab. (Manabe et al., PNAS, 2008) | serum | IHC (1:200) | PFA |
| THBS1 | Mouse | Santa Cruz | sc-59887 | IHC (2 μg/ml) | PFA |
| TNFAIP6 | Rabbit | Sigma | HPA050884 | IHC (10 μg/ml) | PFA |
| TNFAIP6 | Rabbit | Thermo Fisher | PA5-62259 | IHC (2 μg/ml) | PFA |
| vitrin | Rabbit | Novus | NBP2-49453 | IHC (2.5 μg/ml) | PFA |
| VWA1 | Rabbit | Sekiguchi lab. (Manabe et al., PNAS, 2008) | serum | IHC (1:100) | PFA |
| **Secondary** | | | | | |
| Goat IgG Alexa 555 | Donkey | Life Technologies | A21432 | IHC (1:500) | PFA/acetone +acid |
| Guinea pig IgG Alexa 488 | Goat | Life Technologies | A11073 | IHC (1:500) | PFA |
| Hamster IgG Alexa 546 | Goat | Life Technologies | A21111 | IHC (1:500) | PFA |
| Mouse IgG2a Alexa 488 | Goat | Life Technologies | A21131 | IHC (1:500) | PFA |
| Mouse IgM Alexa 488 | Goat | Life Technologies | A21042 | IHC (1:500) | PFA |
| Rabbit IgG Alexa 488 | Goat | Life Technologies | A11034 | IHC (1:500) | PFA/acetone +acid |
| Rabbit IgG Alexa 555 | Goat | Life Technologies | A21429 | IHC (1:500) | PFA/acetone +acid |
| Rat IgG Alexa 488 | Donkey | Life Technologies | A21208 | IHC (1:500) | PFA/acetone +acid |
| Rat IgG Alexa 488 | Goat | Life Technologies | A11006 | IHC (1:500) | PFA |
| Rat IgG Alexa 555 | Goat | Life Technologies | A21434 | IHC (1:500) | PFA/acetone +acid |
| Sheep IgG Alexa 647 | Donkey | Life Technologies | A21448 | IHC (1:500) | PFA |
